# A Lake Charr Pangenome Reveals Highly Conserved Ohnologs as Drivers of Phenotypic Diversity

**DOI:** 10.64898/2026.07.03.729964

**Authors:** Christopher A. Osborne, Nathan J.C. Backenstose, Daniel J. MacGuigan, Steven J. Fleck, Brian F. Lantry, Victor A. Albert, Dimitry Gorsky, Trevor J. Krabbenhoft

## Abstract

Whole-genome duplication (WGD) is hypothesized to spur evolutionary diversification by producing genome-wide duplicate gene sets (Ohnologs) that are initially functionally redundant but can diverge markedly as the effects of relaxed selection accumulate over time. However, the underlying mechanisms remain unclear, in part because genomic studies often reconstruct Ohnolog evolution over millions of years, during which subsequent mutations can obscure deep-time signals. Investigating the relationship between Ohnolog evolution and diversification on a contemporary timescale offers clearer insights. We explore this relationship in Lake Charr (*Salvelinus namaycush*), where ∼10% of genes are retained highly conserved polyploid duplicates following the Salmonid-Specific Fourth Round WGD. Using 31 chromosome-level assemblies of Lake Charr from morphologically and ecologically diverse populations, joined into a pangenome graph, we characterized 189,555 structural variants (SVs) that were significantly less likely to affect genes retained as sequence-conserved Ohnolog pairs, nuancing the hypothesis that gene redundancy, relaxed selection, and functional diversification are intertwined. However, we found that SVs affecting such conserved Ohnologs may be potent drivers of adaptive evolution. Notably, we identified a putative 938-Kb interchromosomal translocation containing 25 genes with highly conserved Ohnologs in a paralogous (but untranslocated) genomic block. This putative translocation appears to have facilitated Ohnolog divergence in *ankrd11* and *hp*, genes putatively linked to craniofacial and lipid metabolic diversity in sympatric Lake Superior morphs. This research reveals that conserved Ohnologs previously presumed to be redundant remain a reservoir for adaptive change.

## INTRODUCTION

Whole-genome duplication (WGD) is hypothesized to drive evolutionary diversification in multiple lineages by generating a reservoir of gene copies, known as Ohnologs^1,2^. In the short term these Ohnologs are redundant copies, but in many systems, both animals and plants^1,3–5^, those duplicates that survive deletion can functionally diverge over time and contribute to adaptation. This arises when one or both gene copies experience relaxed selective pressures, allowing for the development of novel and/or partitioned functions that facilitate biological diversification. However, the relationship between WGD and diversification is complex, and the mechanisms that connect the two are not fully understood. For example, the WGD Radiation Lag-Time Model highlights the often-observed asynchronistic relationship between the timing of a WGD and the timing of species proliferation^6^. The Lineage-specific Ohnologs Resolution Hypothesis proposes that, in some cases, this lag can be explained by a stepwise process of rediploidization (*i.e*., the transition from tetrasomic back to disomic inheritance)^7^. In this model, some regions of the genome rapidly return to a diploid state after WGD and before any major lineage diversification (called Ancestral Ohnolog Resolution events; AORe), while others do so later, after speciation or reproductive isolation has occurred, sometimes after millions of years of apparent stasis^7^. Delayed episodes of post-speciation rediploidization are termed Lineage-specific Ohnolog Resolution events (LORe)^7^. Because LORe (i.e., the second wave of rediploidization) occurs after reproductive isolation, the onset and extent of sequence divergence and fractionation (gene loss) between Ohnologs that rediploidization during this time exhibit lineage-specific patterns^7^ and are hypothesized to be associated with the WGD-induced biological diversification^7,8^. While LORe are hypothesized to facilitate adaptive evolutionary changes^7,8^ it is also possible that these events can be entirely explained by neutral evolutionary forces. For example, genetic drift during speciation-associated bottlenecks (e.g., via founder effects^9^) could fix random mutations in duplicated genes, which could also create lineage-specific patterns of Ohnolog divergence. It remains unclear whether LORe events are associated with adaptive evolutionary divergence or if they are largely the product of neutral evolutionary processes^10^.

Perhaps the best-studied example of the Lineage-specific Ohnologs Resolution hypothesis is the Salmonidae-Specific Fourth Round WGD (SS4R) event that occurred in the last common ancestor of all members of the salmonid family approximately 106 million years ago (Mya)^7,11,12^. Immediately after the SS4R, the ancestral salmonid genome underwent an initial wave of rediploidization, resulting in approximately 60% of the genome reverting to a diploid state^7,11,12^. After approximately 40 My of relative stasis, a second wave of rediploidization occurred, potentially coincident with the split of the three salmonid subfamilies^7,11,12^. Ohnolog pairs exhibit distinct, lineage-specific patterns of sequence divergence and gene loss across salmonid subfamilies, suggesting the second wave of rediploidization may have been the primary driver of diversification observed in Salmoniformes^13^. Ohnologs that began rediploidization during the second wave (hereafter, late-diverging Ohnologs) still exhibit significantly higher levels of sequence similarity, have similar tissue-specific gene expression patterns, and epigenetic marking conservation than those that rediploidized shortly after the SS4R (hereafter early-diverging Ohnologs)^7,10,11,14^. As such, theory predicts that late-diverging Ohnologs should still exhibit high functional redundancy and, therefore, have greater potential to contribute to future intraspecific-diversification^2^. However, while lineage-specific Ohnolog resolution has been associated with salmonid diversification at the subfamily and generic levels, an open question is the extent to which Ohnologs contribute to subsequent cycles of diversification, long after rediploidization.

Salmonids exhibit remarkable intraspecific phenotypic variability and rapid sympatric adaptive radiation^15,16^, a phenomenon strikingly exemplified by charr (*Salvelinus* spp.), which boast as many as 12 reported sympatric morphs^17–22^. Lake Charr (*S. namaycush*), for example, have developed extensive morphological, behavioral, ecological, and physiological diversity during recent radiation events, driven by abundant ecological opportunities in North American post-glacial lakes^17,23^. Understanding how late-diverging Ohnologs influence this intraspecific diversity may shed light on the latent effects of delayed rediploidization on biological diversification and help explain the lag between WGD and subsequent evolutionary radiation.

While advances in sequencing technologies have enabled considerable insight into the genomic basis of diversification across *Salvelinus* spp. and other salmonids^24–28^, most studies have relied on short-read sequencing mapped to a single reference genome. This strategy is poorly suited for detecting genomic structural variants (SVs) – mutations affecting at least 50 base pairs (bp)^29^. Structural variants can play an outsized role in genome evolution^30–32^, including rediploidization^11,33–36^; therefore, studies that accurately characterize SVs are essential to fully understand the continuing influence of the SS4R on diversification. The advent of long-read sequencing has greatly facilitated such investigations by improving genome assembly and enabling the development of pangenomes, which are multi-genome graphs that capture all shared and unique sequences within a population, species, or lineage. These genomic resources provide robust, unbiased scaffolds for detecting SVs^37–40^. While comparisons of single genomes have been useful for the exploration of SVs in salmonids, previous work focused mainly on recent chromosomal inversions^41^, and have overlooked the potential role of SS4R^42^. A pangenome of morphologically and ecologically diverse charr offers valuable insights into the genomic drivers of diversity in this lineage and further illuminate the impact of delayed rediploidization on vertebrate evolution.

Here, we constructed a high-quality pangenome graph for Lake Charr (*S. namaycush*) which exhibits high levels of recently derived intraspecific diversity, thus providing an ideal system in which to examine how delayed rediploidization might shape recent biological diversification. Although the genomic basis of the *S. namaycush* radiation has been examined in several studies^43–46^, the putative roles of SVs and WGD remain unexplored. By constructing a high-quality pangenome from 31 *S. namaycush* individuals representing 10 morphologically and ecologically diverse populations, we accurately characterize the genomic SV landscape and gain new insights into the role of delayed rediploidization in shaping phenotypic diversity. Notably, we identify a 938-kilobase region with elevated sequence and structural variation potentially associated with an interchromosomal translocation. Structural and sequence variation in this region appear to be linked to key morphological and ecological variation observed among sympatric *S. namaycush* morphs from Lake Superior. Collectively, these results establish our pangenome as a valuable resource for studying structural variation, phenotypic diversity, and population management in *S. namaycush*, and provide broader insight into phenotypic evolution within Salmonidae.

## RESULTS AND DISCUSSION

### Glacial Refugial Ancestry and Recent Isolation in Small Lakes Shape *S. namaycush*

#### Population Structure

To investigate the genomic basis of biological diversity in Lake Charr, we collected samples from 10 populations that exhibit broad ecological and morphological variation. These populations include a dwarf morph from the Adirondack Mountains^47^, Finger Lakes populations that exhibit unique spawning behavior and thiamine metabolism^48–51^, and three sympatric morphs from Lake Superior that exhibit considerable morphological, behavioral, and physiological variation^52^. To establish the demographic framework for our pangenomic investigation, we first assessed intra-population diversity and resolved the genetic structure across these groups by generating moderate-coverage whole-genome Illumina short-read sequences for 29 fish (n = 2–3 per population). A principal component analysis of ∼3.2 million single-nucleotide variants (SNVs) revealed that the genetic structure of these populations reflects their glacial refugia during the Last Glacial Maximum (LGM) of the Laurentide Ice Sheet. Individuals from the Laurentian Great Lakes (hereafter Great Lakes), which likely represent populations descended from a mix of Beringian and Mississippian glacial ancestors, separate along the first principal component (PC1) from eastern populations, which are likely descendants of an Atlantic refugial ancestors (**Fig. 1a, 1b**), supporting previous mitochondrial DNA results^53^. Demographic history reconstruction shows that the ancestral populations of each sample reached their lowest estimated effective population size (*N_e_*) during the LGM, approximately 19 – 33 thousand years ago^54^, consistent with Backenstose et al.^23^. This decline was followed by a precipitous increase in *N_e_* after glacial retreat, likely corresponding to rapid population expansions associated with abundant novel habitat availability as the Great Lakes deglaciated (**Fig. 1c).** Collectively, these results provide further support that historical glacial cycles have broadly shaped contemporary patterns of genetic diversity and population structure of *S. namaycush*^23,53,55^.

**Fig. 1:**
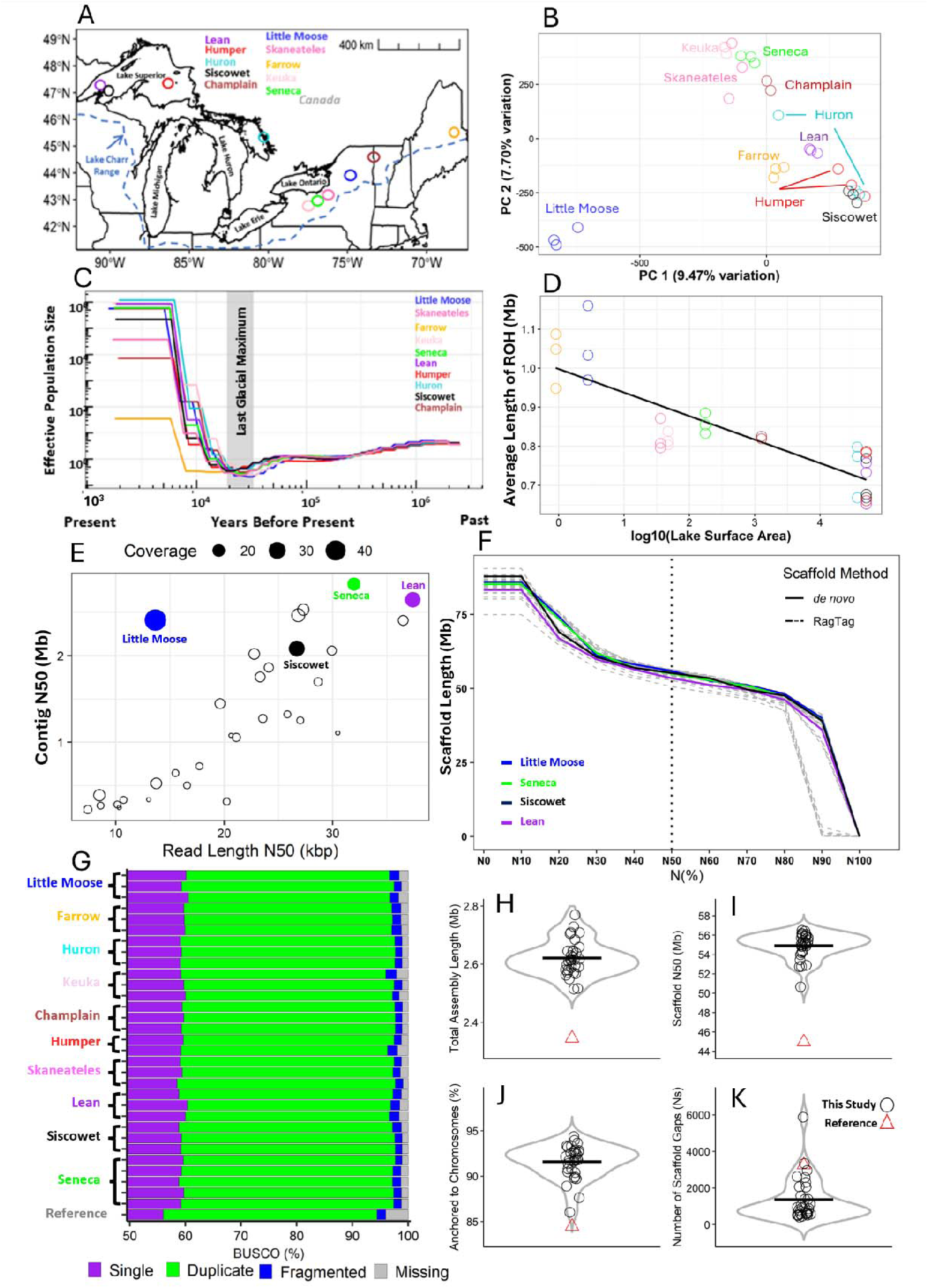
Population structure, demographic history, and genome assembly results of 31 *Salvelinus namaycush*. **A**, Lake of origin for *S. namaycush* used in this study, points are colored according to each lake. **B**, A principal components analysis using genome-wide single-nucleotide variants detected in 10 populations of *S. namaycush*. **C**, Demographic history of *S. namaycush* populations, estimated using the pairwise sequentially Markovian coalescent model (PSMC^158^). The light-grey bar denotes the estimated timing of the last glacial maximum of the Laurentide Ice Sheet, approximately 19 – 33 Kya. **D**, Relationship between the average length of a run of homozygosity and the log10 transformed surface area (in square kilometers) of the lake from which the sample originated. **E**, Relationship between read-length N50 and total estimated genome coverage with the resultant contig length N50 of the assembly produced. **F**, Assembly contiguity, measured by the Nx^th^ length, for all 31 *S. namaycush* assemblies produced in this study. *De novo* assemblies (those scaffolded with Hi-C data) are shown in solid lines and colored according to the lake of origin, and broken lines represent the RagTag scaffolded assemblies. **G**, Benchmark Universal Single Copy (BUSCO) score results for each assembly produced in this study and the current *S. namaycush* reference genome. **H – K**, Violin plots for **H,** total assembly length in megabases (Mb), **I**, scaffold N50 (Mb), **J**, percent of the total assembly length anchored to one of 42 chromosomes, and **K**, the number of scaffolding gaps (denoted as Ns in the assembly) for each new assembly and the current reference genome for *S. namaycush*.

Lake size also had a considerable impact on contemporary genetic diversity and population structure in *S. namaycush*. The Little Moose Lake population - a small lake (2.3 km^2^ surface area) in the Adirondack Mountains of New York - exhibit considerable sequence divergence from all other populations, including the geographically proximate populations in the Finger Lakes (**Fig. 1b)**. The population from Farrow Lake, a small mountain lake (0.9 km² surface area) in Maine, USA, exhibits similar patterns of divergence, as indicated by separation along PC3 (**Fig. S1**), although recent stocking-induced admixture with individuals from Lake Superior (**Fig. S2)** obscures this pattern on the first two principal components. Elevated sequence divergence across short geographic distributions is consistently observed in *S. namaycush* populations occupying small, isolated lakes^56–59^, including other Adirondack populations^60,61^. The average lengths of runs of homozygosity, a proxy for the recency of inbreeding events^62^, strongly correlate with the size of the lake currently occupied by each population (**Fig. 1d),** suggesting the pattern of elevated sequence divergence is likely driven by reproductive isolation and genetic drift in the small, remote lakes. The elevated levels of sequence divergence observed between these small populations and nearby populations support the Drift Barrier Hypothesis^63^, which proposes that small *N_e_* can lead to increased substitution rates, likely due to a hindrance of natural selection’s purging of mutations, allowing for drift to play an outsized role in driving mutations to fixation. The unique genomic characteristics of *S. namaycush* in small, isolated lakes suggest the possibility of elevated endemism, which could be essential for informing future conservation and management decisions for these populations^64^.

### Chromosome-level Assembly and Annotation of 31 *S. namaycush* Genomes

To facilitate whole-genome comparisons, we generated two to five genome assemblies for each population, including four *de novo* chromosome-level assemblies that incorporated high-coverage (>30x) Oxford Nanopore Technology (ONT) long-read sequence data and high-throughput chromatin conformation capture (Hi-C), and 27 genome assemblies with moderate-coverage (15 – 20x) ONT long-reads. We optimized parameters in Flye ^65^ v2.9.3 for each assembly to maximize contig length and assembly completeness without compromising accuracy (**see Methods**). This revealed that read-length N50 had a greater impact on assembly contiguity than genome sequencing depth (**Fig. 1e)**. High-coverage assemblies were scaffolded into chromosomes using Hi-C data, and we implemented a nuanced, reference-guided scaffolding approach for our moderate-coverage assemblies. Briefly, while most studies use a single, highly-contiguous assembly for reference-guided scaffolding, we scaffolded each moderate-coverage assembly using the high-coverage chromosome-level assembly most genetically similar to it, as determined by SNV-based PCA and degree of shared ancestry from an admixture analysis (**Supplementary Table 1, Fig. S1, S2**). This approach led to a comparably high level of contiguity and completeness for our moderate-coverage and high-coverage assemblies (**Fig. 1f, g; see Supplementary Data S1 for full assembly statistics)**.

Our final dataset consisted of 31 high-quality, chromosome-level genome assemblies of *S. namaycush,* each of which was an improvement over the current reference genome for this species across all but one assembly quality metric for two samples (**Fig. 1h – k)**. These improvements included an average increase of 200 Mb in total assembly length (**Fig. 1h**) while also increasing the proportion of each genome anchored to chromosomes by an average of 8% (**Fig. 1j**), resulting in an average increase in scaffold N50 of 10.2 Mb (**Fig. 1i**). Utilizing our nuanced, reference-guided scaffolding approach likely reduced inter-population reference bias, and thus the genome assemblies we produced more accurately represented the full genetic diversity of the populations^66,67^.

We used a combination of *ab initio* and homology-based gene prediction methods using publicly available RNA sequencing data for *S. namaycush* and protein evidence from several publicly available salmonid genomes to predict gene models for each new assembly (**Supplementary Table 2**). Total gene counts in our annotations ranged from 53,511 to 57,544, with an average of 55,045. Complete Benchmark Universal Single Copy Orthologs (BUSCO) scores ranged from 94.3% to 96.0%, with an average of 95.5%, and duplicated BUSCO scores ranged from 36.1% to 39.4% with an average of 37.8, indicating a high degree of completeness in our annotations (see **Supplementary Table 3** for full annotation statistics).

These chromosome-level assemblies now capture the breadth of genomic structure present across *S. namaycush* populations, providing a far more complete view of this species’ genetic landscape than previously possible. With this foundation, questions about the origins of morphological divergence and the genomic basis of ecological specialization can be addressed at a resolution previously out of reach.

### A High-Quality Pangenome Reveals the Structural Variation Landscape of *S. namaycush*

To characterize the genomic structural variation landscape of *S. namaycush*, we constructed a pangenome graph with the 31 newly assembled genomes using the Pangenome Graph Builder pipeline (PGGB^68^ v0.5.4). To ensure that only high-quality SVs were retained, we employed two additional SV detection methods SVIM-ASM^69^ and Sniffles2^70^, and retained only those SVs identified by at least two of the three methods. This resulted in a final dataset of 189,555 high-confidence, non-redundant SVs, including 99,054 (52.26%) deletions, 89,308 (47.11%) insertions, 856 (0.45%) duplications, and 337 (0.18%) inversions (**Fig. 2a, b)**. The average number of SVs detected per assembly was 58,501, ranging from 46,629 to 67,448 (**Fig. 2a, b).**

**Fig. 2:**
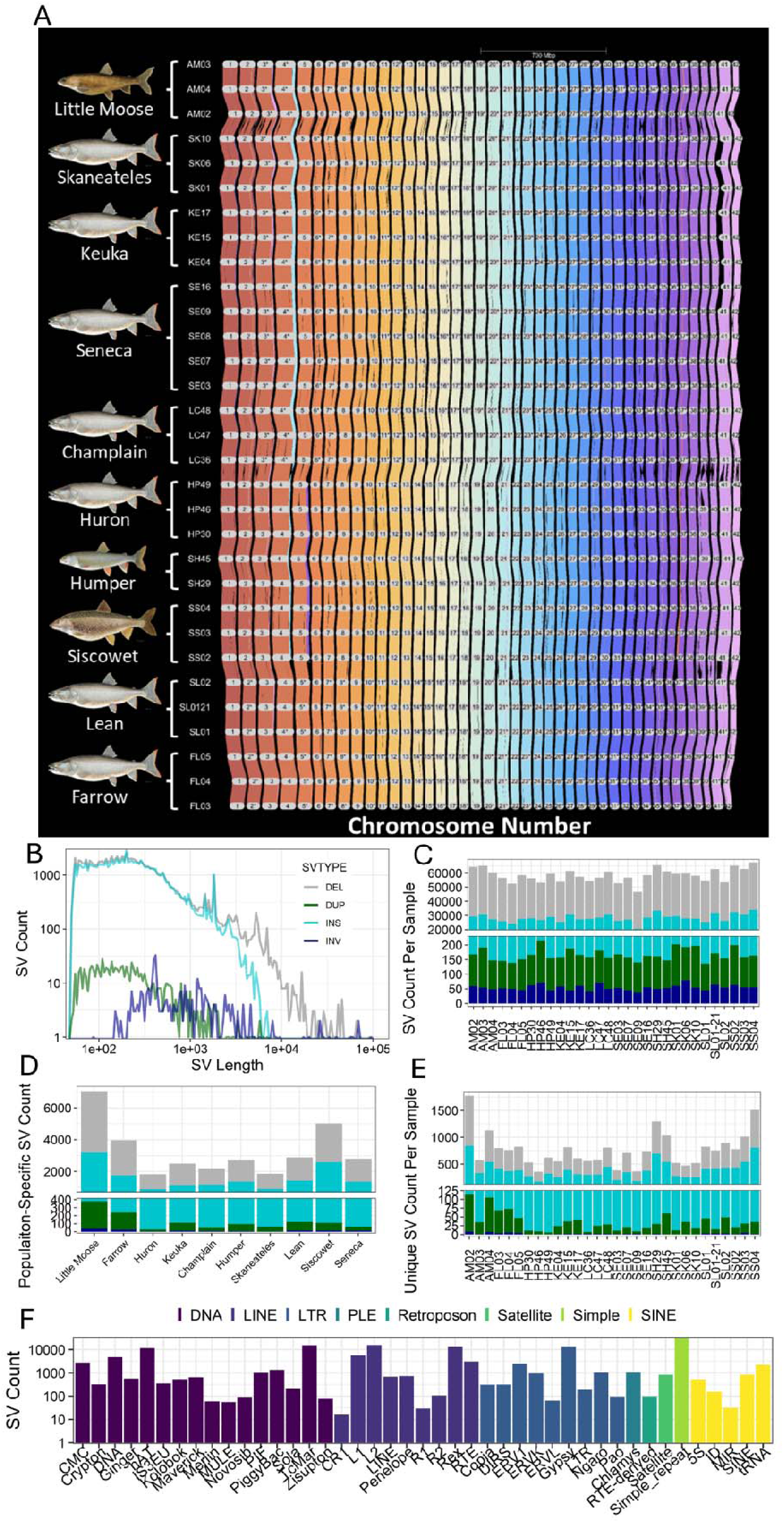
Genomic Structural Variation in *Salvelinus namaycush*. A riparian plot depicting conserved synteny and structural variations in the 42 chromosomes of *S. namaycush* genome. Bands are colored according to which chromosomes the genes of a syntenic block are found in the Farrow Lake genome assembly, which was arbitrarily chosen for visualization purposes. **B**, A frequency plot of the length profile of deletions (DEL), duplications (DUP), insertions (INS), and inversions (INV), the y-axis is log10 transformed. **C**, The total number and type of SVs found in each sample. SV types are colored according to the color legend in plot **B**. **D,** Number and type of SVs found exclusively in each population. **E**, SVs found uniquely in a given genome assembly. SV types in plots **B**-**E** are colored according to the color legend in plot **B**. **F**, SV counts found to overlap different types of repeat elements.

Most pangenome studies use a single high-quality assembly from each population to construct pangenome graphs and then genotype a larger set of individuals using read-to-graph alignment methods^71–73^. While this approach clearly facilitates drastic improvements in the study of SVs, only incorporating a single genome assembly from each population in the SV discovery process (i.e., pangenome construction) likely leads to intrapopulation-level reference bias. By implementing an optimized genome assembly method for our moderate-coverage assemblies, we were able to incorporate multiple high-quality genomes from each population into our SV discovery pipeline without drastically increasing the cost of our study. This approach led to the discovery of many SVs that might have been missed if any single assembly had been used to represent a given population; instead, all individuals were incorporated into a far more variant-rich pangenome **(Fig. 2c**). Of the 32,809 population-specific SVs (SVs found in at least one individual from a given population and absent from other populations; **Fig. 2d**), 19,121 (58.2%) were uniquely detected in only one assembly (**Fig. 2e**). These results highlight the benefit of our moderate-coverage assembly and nuanced scaffolding approaches; a subset of population-specific SVs would have gone undetected if a single individual from each population had been used as the sole reference genome for that population during pangenome construction.

### Association Between Sequence Repeats and SVs

To investigate the association between SVs and transposable elements in the *S. namaycush* pangenome, we identified the type and frequency of repeat elements overlapped by insertions and deletions (**Fig. 2f**). DNA transposons were the most common repeat class associated with SVs, with the TC1 superfamily of repeats being the most pronounced among them. Simple repeats were the most common single category of repeat type associated with SVs. The association of TC1 repeats with SVs is unsurprising considering TC1-Mairner repeats are the most common repeat type in the *S. namaycush* genomes assembled here (10.6 – 12.1% of the total genome length, a proportion consistent with other salmonids^11,74^). The high frequency of SVs associated with TC1-mariner suggests recent activity of these transposable elements, as observed in the Rainbow Trout (*Oncorhynchus mykiss*) genome^75^. Alternatively, the high abundance of TC1-mariner repeats in salmonid genomes might cause them to serve as nucleation points for homologous recombination, leading to structural variants induced by unbalanced recombination or non-allelic homologous recombination, as has been suggested for Alu repeats in the human genome^76^. However, a recent assessment of recombination hotspots in multiple salmonid genomes showed no significant correlation between TC1-mariner repeats and recombination frequency^77^, suggesting that TC1-mariner transposition is the more likely explanation for the pattern observed here. Overall, the association observed between TC1-mariner repeats and SVs suggests that TE activity may remain a dominant generator of structural variation in *S. namaycush*. In fact, persistent TE-driven mutability may, where it exists, contribute to genomic fluidity^11,78^ and has been associated with divergence among Ohnologs in salmonids^74,79^. As such, TE dynamics may represent an important mediator of salmonid genome evolution that should be considered in further studies, alongside the influence of WGDs.

### Rediploidization Age Affects Structural Variation Patterns

To investigate the role of delayed rediploidization in shaping the SV landscape in *S. namaycush*, we first identified early- and late-diverging syntenic blocks (**Methods**), revealing 9,333 early- and 2,316 late-diverging Ohnolog pairs (**Fig. 3a)**. Mapping these Ohnolog pairs to their genomic location revealed that late-diverging Ohnologs are largely contained in 16 regions (i.e., eight homeologous pairs) concentrated near the telomeric ends of chromosomes (**Fig. S3)**, consistent with previous assessments in Atlantic Salmon (*Salmo salar*)^11,80^. In support of the greater functional redundancy of late-diverging Ohnologs, we show that CpG DNA methylation (i.e., 5-methylcytosine) patterns are significantly more similar between late- than early-diverging Ohnologs (Wilcoxon rank-sum test, *p* < 0.001; **Fig. 3b**). A greater degree of similarity in the epigenetic regulation of late- vs early-diverging Ohnologs has also been documented in patterns of histone H3 lysine 27 acetylation (H3K27ac) peaks in Atlantic salmon^14^ and is congruent with a higher degree of similarity in gene expression patterns^7,13^.

**Fig. 3:**
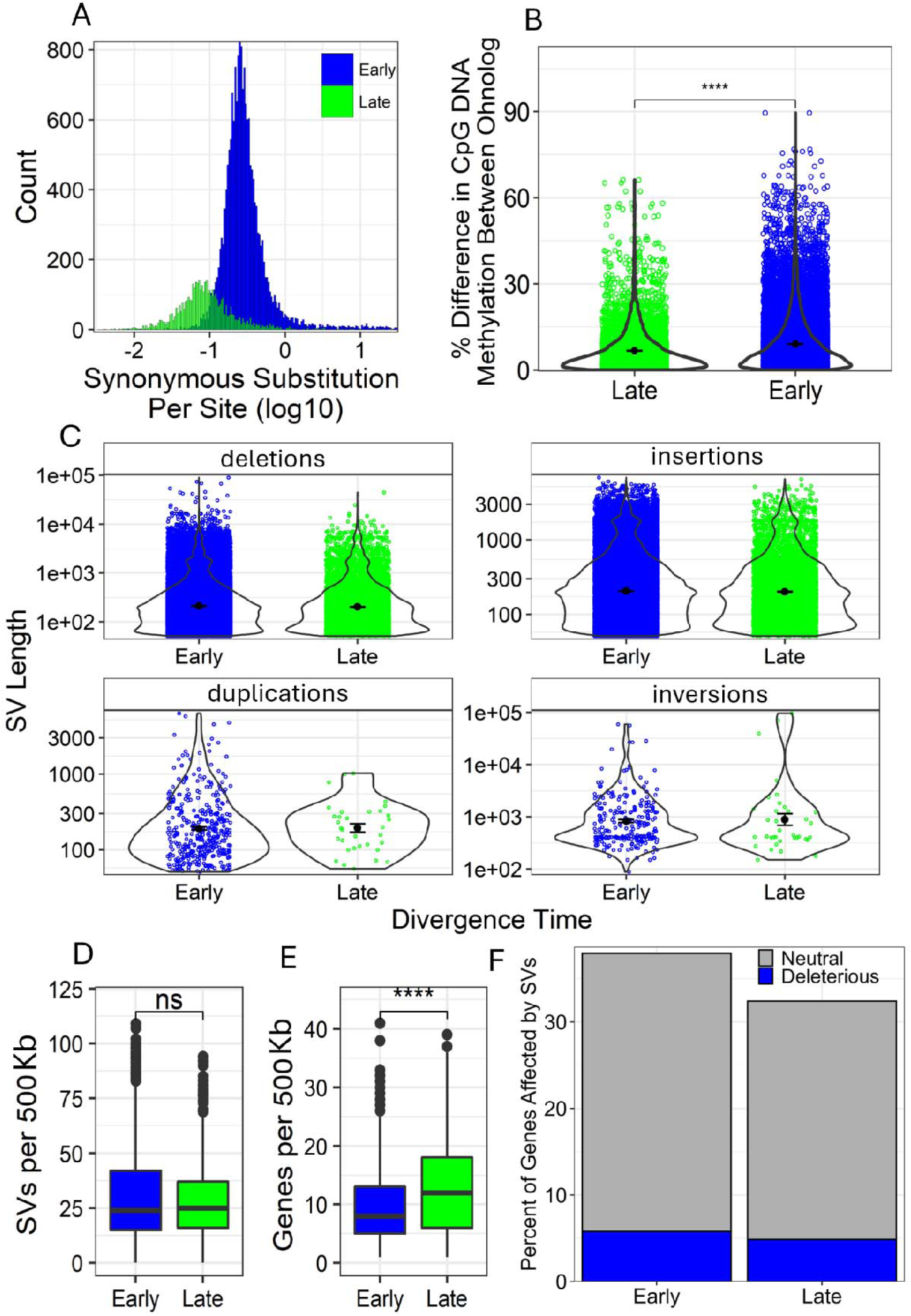
Structural Variation Characteristics of Early- and Late-diverging Genomic Regions. **A**, Histogram of synonymous substitutions per site for early- and late-diverging Ohnologs. **B**, Violin plot showing the percent difference in 5-methylcytosine CpG DNA methylation between early and late-diverging Ohnologs. Significant differences determined using Wilcoxon rank-sum tests (**** *p*< 0.001) are shown. **C**, Violin plots showing lengths of deletions, insertions, duplications, and inversions found in early- and late-diverging regions. For all violin plots, black dots represent means, and horizontal error bars represent the standard error. **D** and **E**, Box plots comparing SV and gene densities (number per 500 Kb window) from early- and late-diverging syntenic blocks. Statistical significance, evaluated using Chi-Squared Tests, is displayed (**** *p*< 0.001); **ns** denotes no significant difference was detected. **F**, Bar chart of the percent of genes from each region affected by SVs, blue bars represent the proportion of genes affected by putatively deleterious SVs (i.e., SVs overlapping exons and promoters).

The average length (**Fig. 3c**) and density (number of SVs per 500 Kb window; **Fig. 3d)** of SVs were similar between early- and late-diverging regions (Wilcoxon rank-sum tests, *p* > 0.05). In contrast, Bertolotii et al.^80^, found SVs were significantly less common in late-diverging regions of the Atlantic Salmon genome. This difference is likely due to short-read sequencing (SRS) data being used to detect SVs, as late-diverging regions had erroneously inflated read depths (defined by the authors as >10x higher than the genome-wide average), resulting in SVs detected in these regions being more likely to be excluded from further analyses due to concomitantly high false discovery rates^80^. The difficulty of mapping SRS data to late-diverging regions also leads to high error rates in SNV detection^81^, and in identifying recombination markers in a linkage map^82^ in other salmonids. While we also observed this with our SRS data, we found this problem was substantially reduced when mapping long-read data to late-diverging regions (**Fig. S4**). Long reads are more likely to span larger sections of homeologous regions and may even extend into flanking unique sequences, similar to how long reads facilitate improved detection of various types of SVs^83,84^. This improved read-mapping accuracy likely provides the necessary anchoring information to distinguish true mapping locations from homeologous regions. These results highlight the value of long-read sequencing for investigating complex genomic phenomena in organisms that have undergone recent polyploidy.

Of the 44,396 SVs overlapping genes (23.42% of all SVs), 22,461 affected early-diverging Ohnologs, 4,112 affected late-diverging Ohnologs, and 17,823 affected non-Ohnologous genes. Although gene density was significantly higher in late-diverging regions (Wilcoxon rank-sum test, *p* < 0.001; **Fig. 3e**), a significantly smaller percentage of late-diverging Ohnologs were affected by SVs (32.8%, n=1501) than early-diverging Ohnologs (37.9%, n=7079; X^2^ = 77.661, *df* = 1, *p* < 0.001; **Fig. 3f**). We also observed a marginally higher percentage of genes with putatively deleterious SVs (i.e., those affecting exons or promoters) in early-diverging (5.8%, n=1081) than in late-diverging Ohnologs (4.9%, n=225), although this difference was not statistically significant (**Fig. 3f**). Given that the length and density of SVs were similar between regions, while gene density was significantly higher in late-diverging regions, one might expect a larger proportion of late-diverging genes to be affected by SVs, yet we observe the opposite pattern. This appears to contradict what theory predicts about relaxed selection on redundant genes. That is, given that late-diverging Ohnologs are far more similar, both genetically and epigenetically, they should be more tolerant to putatively deleterious mutations, as disruption of one gene copy should be buffered by the retained functionality of its counterpart ^2^. Furthermore, recombination rates are significantly elevated in late-diverging regions in salmonids, likely due to higher copy numbers of PRDM9-binding sites^77^, also suggesting that SV density should be elevated in these regions, given the strong correlation between recombination hotspots and SV accumulation^85,86^. Instead, the lower frequency of SVs affecting late-diverging Ohnologs suggests these genes might be less tolerant to SV-induced perturbations. Retention of high sequence similarity between late-diverging Ohnologs might be driven by selective pressures on gene-product stoichiometry (i.e., gene dosage)^10,14,87,88^, but this phenomenon has not been tested at the intraspecific level. Future studies that couple pangenomes with multi-tissue gene expression data would likely represent a robust means of testing this hypothesis.

While a mechanistic explanation for the maintenance of sequence similarity between late-diverging Ohnologs is beyond the scope of this study, Allendorf et al.^89^ provided a theoretical framework for how periodic recombination between Ohnologs might lead to gene conversion events that purge mutations from these genes. Multivalent pairing leading to recombination events between late-diverging genomic regions is still observed in many salmonid species^90–92^, including *S. namaycush*^93^, and can result in gene conversion events^94,95^. Although recombination between late-diverging regions appears to occur primarily in males ^91^ and may be infrequent^96^, gene conversion events might only need to occur approximately once per generation to ensure that even neutral alleles are preserved in a population for long periods^97^.

Viewed together, our results are consistent with the hypothesis that late-diverging Ohnologs remain embedded in dosage-sensitive regulatory architectures that limit the accumulation of structural variants. Alternatively, recurrent gene conversion between homeologous regions may periodically reset divergence and maintain functional similarity. Both mechanisms would generate the unexpected pattern observed: high sequence similarity but low SV tolerance in late-diverging regions. This observation raises the larger question of whether rediploidization age predicts the genomic components most available for evolutionary experimentation, a hypothesis that can be addressed in future work by integrating our pangenome with gene expression and chromatin-based data.

### Late-Diverging Ohnologs are Involved in Phenotypic Diversity in *S. namaycush*

*Salvelinus namaycush* exhibits an impressive degree of phenotypic variation across its range; morphs differ widely in color, body size and shape, lipid composition and physiology, dietary habits, partial anadromy, and phenology, with several instances of multiple morphs occurring in sympatry^16,98–101^ (reviewed in^17,102^). Indeed, historical records report as many as 12 sympatric morphs in Lake Superior^22^, before anthropogenic stressors led to widespread local extirpation events^103^. Notably, the phenotypic variation exhibited by *S. namaycush* appears to represent a recent radiation event, as this species has primarily diversified across depth gradients and through resource and niche partitioning as the Canadian and Laurentian Great Lakes deglaciated and concomitant ecological opportunities arose^23,98,104,105^. While the biology and ecology of many of these morphs have been well studied, the genomic basis for this broad phenotypic variation has only been explored using methods ill-equipped to detect SVs, such as microsatellites^59,105–112^, mitochondrial genomes^53,113,114^, transcriptomes (RNA-seq)^45,115,116^, restriction site-associated DNA sequencing (RAD-seq)^43,57,117,118^, and only very recently, using low-coverage whole-genome sequencing^46^. Although these studies have demonstrated a heritable genetic basis for some of the phenotypic diversity in *S. namaycush*, the inherent limitations in the approaches used have provided little insight into the genomic architecture of this radiation.

To investigate the putative role of late-diverging Ohnologs on shaping phenotypic diversity in *S. namaycush*, we use our pangenome to compare the genomes of three extant sympatric morphs from Lake Superior: lean, siscowet, and humper. These morphs are primarily distinguished by their habitat selection, lipid physiology, and morphology^22,52,119–122^ (**Fig. 4**). Our pangenome analysis revealed a non-reciprocal (i.e., insertional) translocation of a 938 Kb segment of chromosome 1 to chromosome 3 in the assemblies of two of the three siscowet and both humper samples (**Fig. 5a, Fig. S5**). Interestingly, the translocated segment of chromosome 1 is a late-diverging region that was moved to an early-diverging region of chromosome 3 (**Fig. 5a**). To improve our understanding of this translocation, we used haplotype-aware error correction of the ONT long-read and Hi-C data for the high-coverage lean and siscowet samples to generate haplotype-resolved genome assemblies for each. Mapping each haplotype to the other revealed that the siscowet was heterozygous for the translocation, while the translocation was absent from both lean haplotypes (**Fig. 5a**). A closer inspection of the long-read alignments at this locus revealed that the translocation breakpoint was only supported by two ultra-long (>72 kb) reads, while all other reads mapped very poorly to the region (Fig. S6). The fact that only these two reads span the breakpoint suggests this may have been an assembly error due to a physical chimeric molecule generated during the ONT ligation process. These reads are very similar in length and map to identical coordinates in reverse complement, suggesting they may represent an Oxford Nanopore read duplex—the template and complement strands of a single, rather than two independent biological chimeras. Because assemblers like Flye aggressively prioritize ultra-long reads to resolve complex graphs, this single duplex molecule either provided the only strong support for what is a real translocation or forced an artificial path during contig layout. The heterozygous state of this translocation in siscowets likely explains why the translocation was missing from one of the three assemblies for this morph. That is, in one of the unphased assemblies, the chromosome 1 and 3 haplotypes without the translocation were retained by chance.

**Fig. 4.**
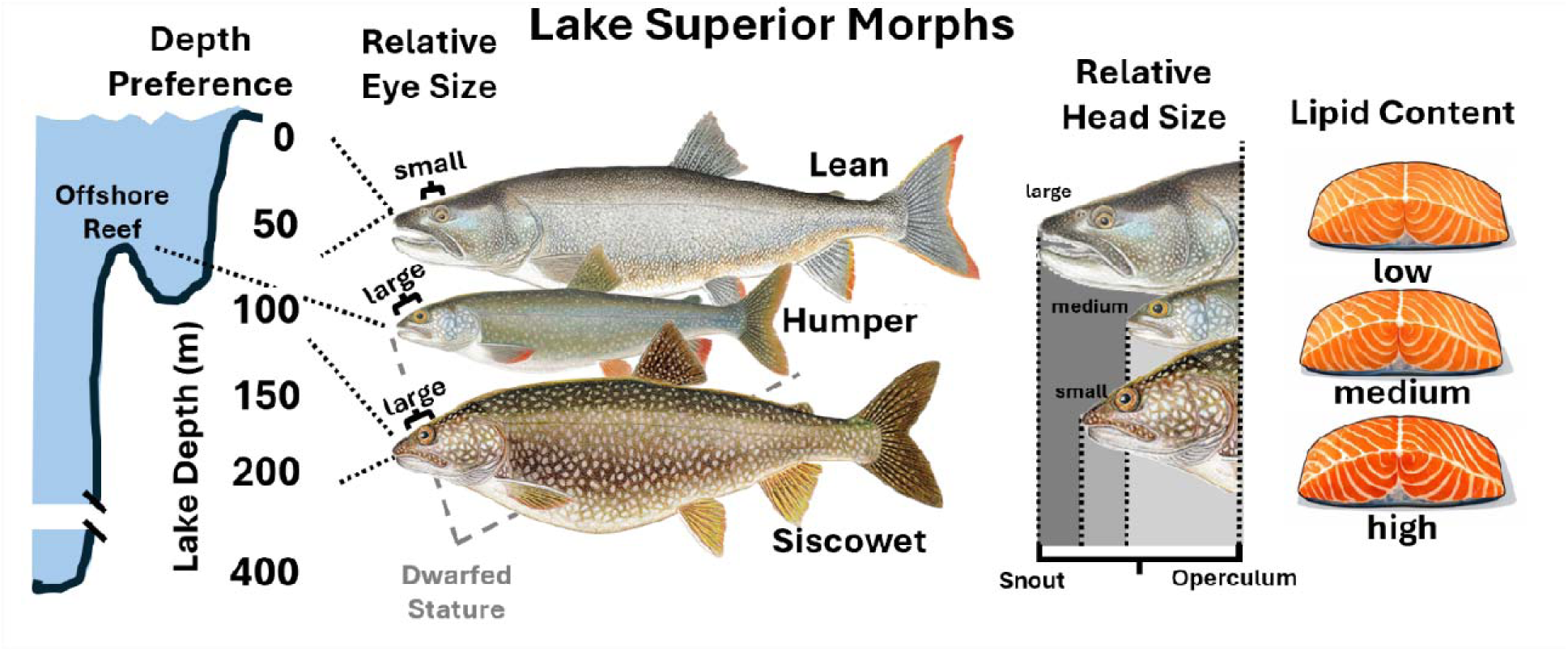
Phenotypic variation among Lake Superior Lake Charr (*Salvelinus namaycush*) morphs. A diagram depicting the morphological and physiological traits of three *S. namaycush* morphs—lean, humper, and siscowet—relative to their ecological niches. **(Left)** Depth preferences correlate with eye size and body stature; deeper-dwelling morphs (humper and siscowet) exhibit larger relative eye sizes and the humper has a small body size compared to the shallow-water lean morph. **(Center)** Variation in relative head size, measured from snout to posterior extent of operculum, distinguishes the morphotypes, with the lean morph possessing the largest relative head size. **(Right)** Cross-sections of muscle tissue indicate varying lipid content, showing an increase in fat storage from lean (low) to siscowet (high), an adaptation likely related to buoyancy regulation in deepwater environments.

**Fig. 5:**
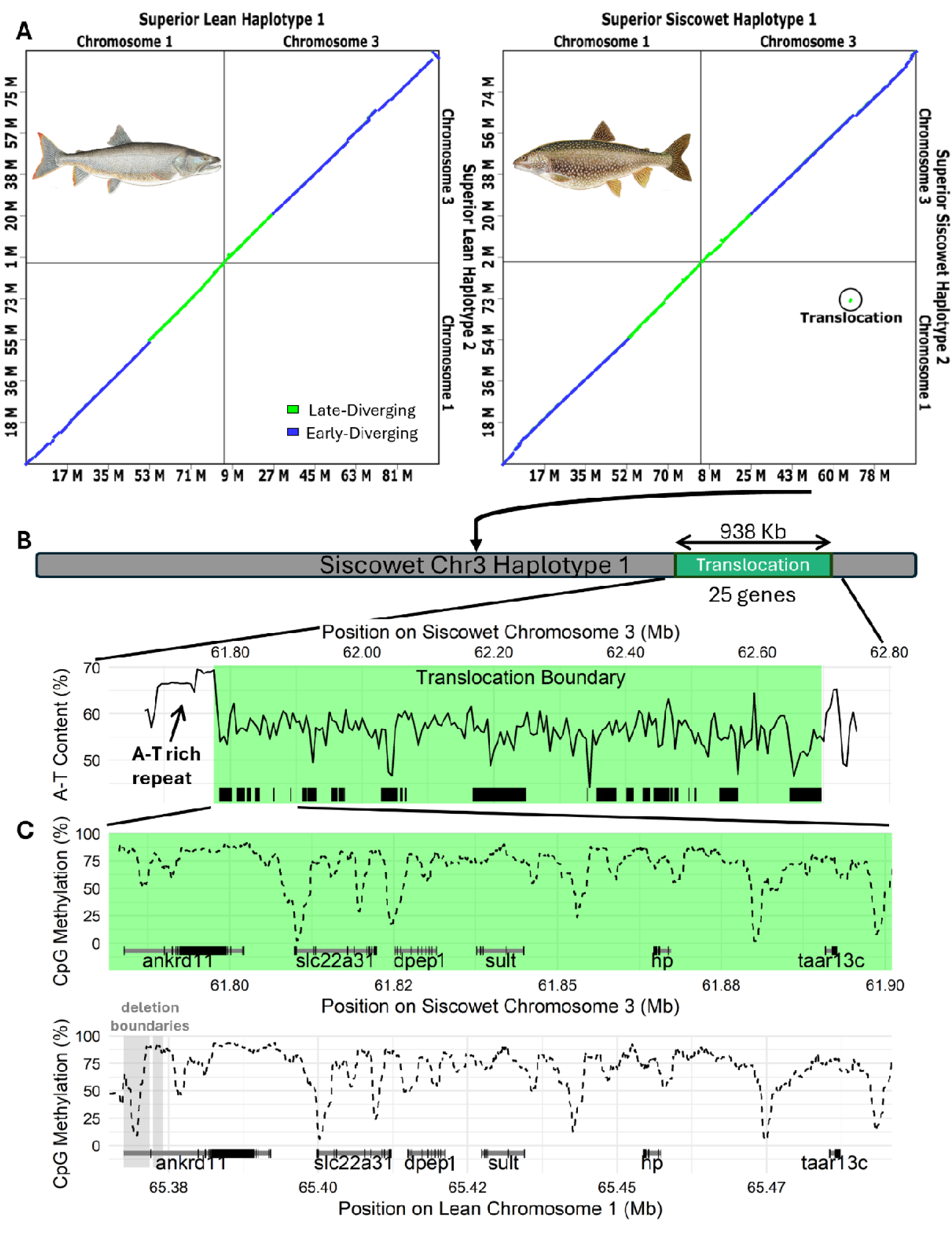
Putative interchromosomal translocation discovered in Siscowet Morph. **A**, Dot plot depicting the sequence alignment of haplotype resolved chromosomes 1 and 3 in lean (left) and siscowet (right). Synteny between siscowet chromosome 1 from haplotype 2 and chromosome 3 from haplotype 1 indicates it is heterozygous for an interchromosomal translocation between these chromosomes. **B**, Percent A-T content of the translocation occurred and its flanking regions in siscowet. A-T content was calculated in 50 Kb non-overlapping windows. A green box depicts the boundary of the translocation, and the start and end positions of the 25 genes found on the translocation are displayed as black boxes. **C**, A comparison of 5-methylcytosine CpG DNA methylation and gene structure of the first five genes on the translocated segment in siscowet chromosome 3 and the untranslated homologous region from lean chromosome 1. The dashed line represents the rolling average of percent methylation calculated for 20 CpG dinucleotide windows. The exons (black boxes) and introns (grey thin bars) are displayed for each gene in the region. The boundaries of two deletions affecting *ankrd11* in siscowet morphs are depicted as grey vertical boxes.

Regardless, the underlying genomic architecture appears to be a hotspot for morph-specific divergence. A recent study used high-density RAD-seq data to explore genomic regions of divergence among lean, siscowet, and humper morphs of *S. namaycush* from the same Lake Superior populations used here^123^. That study found the region of the genome that most strongly distinguished between the morphs occurred between 60 and 80 Mb on chromosome 1. A synteny analysis with the reference genome used in that study revealed that the putative translocation we found here falls squarely within that region (**Fig. 7a**). Moreover, three of the SNVs they found to most strongly differentiate lean from siscowet and humper morphs occurred almost exactly at the midpoint of the translocation (**Fig. S7b**). The RAD-seq approach implemented by the previous study identified the same region as being highly divergent between morphs, and the authors of that study believed the region contained a large chromosomal inversion that was polymorphic among the morphs^123^. While we were not able to definitively resolve the genomic architecture of this region, the concentration of outlier SNVs and structural divergence suggest a localized reduction in recombination, driving concomitant sequence divergence that may play a role in maintaining the phenotypic segregation between deepwater and shallow-water morphs. We discovered a large ∼15 Kb AT-rich repeat region at one of the boundaries of the translocation on chromosome 3 in siscowet (**Fig. 5b).** We hypothesize that this AT-rich repeat may have facilitated an interchromosomal translocation, as AT-rich repeats, particularly long palindromic repeats, are the most common source of non-Robertsonian translocation in humans^124^, such as the recurrent 11q22 interchromosomal translocation associated with Derivative 22 (der22) syndrome^125^. While the AT-rich region found here did not contain large palindromic regions, it is possible that after causing the translocation, likely in the last common ancestor of siscowet and humpers, mutations in the AT-rich repeat region may have degraded what was once a palindromic repeat, effectively stabilizing the region. Akgun et al.^126^ proposed that the accumulation of mutations that would disrupt such palindromic sequences would be favored by natural selection, as they would stabilize such regions and prevent further mutations. This is especially likely if the translocation conferred a selective advantage on the carrier.

To investigate additional genomic divergence in this region, we identified the gene content and compared coding sequences and epigenetic characteristics between the leans and siscowet. Twenty-five genes were in this region (**Fig. 5b**, see **Supplementary Table 4** for the complete gene list), two of which exhibited notable epigenetic or structural polymorphism between morphs. Notably, we found two deletions in siscowet affecting the ankyrin repeat domain-containing protein 11 (*ankrd11*), one of which appears to have disrupted a hypomethylated valley present in the first intron of this gene (**Fig. 5c, Fig. S8**). Remarkably, in humans, a homologous deletion affecting an enhancer in the first intron of *ankrd11* causes KBG syndrome in carriers, a disorder characterized by brachycephaly, macrodontia, hand size abnormalities, and, in extreme cases, shortened stature^127–130^. These conditions accurately describe the craniofacial and limb morphological characteristics of both siscowet and humper morphs relative to the lean morph^52^. Consequently, we hypothesize a conserved role for *ankrd11* across vertebrates and implicate these deletions in the divergent craniofacial and fin morphologies of siscowet and humper morphs. While these results represent the first putative genomic basis of craniofacial variation in *S. namaycush*, similar morphological diversity has been extensively studied in the Arctic Charr (*S. alpinus*)^131–134^. However, thus far, *ankrd11* has not been linked to craniofacial morphology in *S. alpinus,* suggesting a lack of convergent evolution with *S. namaycush*.

Siscowet have evolved to regulate their buoyancy primarily through drastically elevated tissue lipid content, which makes them positively buoyant, while other morphs primarily regulate buoyancy using their gas bladders^135,136^. This change appears to be an adaptation that allows siscowets to undergo large-scale diel vertical migrations from depths exceeding 150 meters (m) to less than 50m in a short period to track prey^104^. This feat could lead to barotrauma in fish that primarily rely on their gas bladders for buoyancy regulation^137,138^. The high lipid content of siscowets is associated with increased lipid sequestration into muscle and liver adipose tissue, yet significantly lower serum lipid levels than leans^115^. In a common garden rearing experiment, Goetz et al.^45^ demonstrated significantly elevated expression of haptoglobin (*hp*) in siscowet compared to leans. Haptoglobin is essential for mitigating oxidative stress induced by free heme in the bloodstream and is significantly upregulated in humans with elevated lipid levels, potentially reducing lipid-induced inflammation^139^. Remarkably, *hp* was also located within the this region of elevated divergence (**Fig. 5c)**. Comparisons with leans revealed the copy of *hp* from this region in siscowet has a 319 bp deletion affecting the third intron and exhibits elevated sequence divergence, including three non-synonymous substitutions. A recurrent deletion in humans increases the antioxidant activity of *hp*, which has been associated with reduced serum lipid levels and, concomitantly, reduces lipid-induced oxidative stress^140^, similar to what has been observed in siscowets. If the mutations we documented in *hp* in siscowet are associated with differential function or expression levels in this morph, they may mitigate lipid-induced oxidative stress associated with their elevated fat content and may have facilitated their divergence into a deepwater morph.

*Salvelinus namaycush*, like other salmonids have been shown to use their olfactory senses to navigate to their natal spawning sites. Specifically, amino acids produced from male salmonids are detected by a suite of trace amine-associated receptor (*taar*) genes, which appear to have experienced a gene family expansion in salmonids. Interestingly, we found that the putative translocation also contained trace amine-associated receptor 13c (*taar13c;* **Fig. 5c**), which indeed plays a role in olfactory reception and response to odorant detection in zebrafish, and has recently been implicated in variations in maturation timing and natal homing in *S. salar*. Although *taar13c* did not exhibit coding sequence or epigenetic differences between morphs, the ins(3;1) translocation might alter its regulation via altering its proximity to cis- and trans-regulatory elements. If this is the case, then the presence of *taar13c* in the translocation could have played a role in shaping spawning site and maturation timing differences observed between these morphs, facilitating partial reproductive isolation^119,141–151^Together, these findings suggest that late-diverging Ohnolog regions can act as focal points for large-effect structural mutations capable of reshaping key ecological and morphological traits. This raises the broader possibility that the genomic legacy of delayed rediploidization continues to channel the evolutionary trajectories available during salmonid radiations. Future studies could test this hypothesis by incorporating gene expression data for each morph collected across the spawning window in *S. namaycush*.

By constructing a pangenome for *S. namaycush*, we provide novel insights into the dynamics of rediploidization and the role of structural variation in shaping biological diversification in a member of the salmon family. We found that, despite late-diverging Ohnologs having significantly higher sequence and epigenetic similarity, and therefore greater redundancy, they were significantly less likely to be affected by SVs than the more differentiated early-diverging Ohnologs. However, we find evidence that, although less common, SVs leading to increased divergence among late-diverging Ohnologs may still play an important role in evolutionary diversification in this species. The large putative interchromosomal translocation potentially linked to morph diversity in Lake Superior contained 25 genes with late-diverging Ohnologs. Notably, some of the genes in this region were affected by additional SVs not observed in other populations. This suggests that a translocation may have played a role in facilitating divergence among these otherwise highly conserved late-diverging Ohnologs, providing an example of how delayed rediploidization might still be affecting biological diversification in *S. namaycush*. The involvement of a late-diverging Ohnolog region in this region indicates that residual WGD architecture can influence where impactful structural variants arise. By connecting deep genome history with recent morphological diversification, these findings highlight a genomic route through which salmonids may repeatedly generate novel ecological forms. The pangenome graph we present broadens our understanding of genome evolution following WGD in salmonids and other vertebrates. These results provide a robust basis for dissecting the origins of this group’s broad phenotypic diversity and for guiding conservation and management strategies for this ecologically, economically, and culturally important species.

## METHODS

### Sample Collection and Morph Identification

Gill nets were used to collect *Salvelinus namaycush* from Lake Superior, Lake Ontario, Seneca Lake, Keuka Lake, Skaneateles Lake, and Farrow Lake (see Supplementary data for full collection location data). Lake Ontario samples were checked for hatchery-implanted coded-wire-tags using a Handheld Wand Coded Wire Tag Detector (Northwest Marine Technology Inc.). Tagged Lake Trout had their snouts removed for recovery of coded-wire tags. Coded-wire-tag codes were read by experienced readers and verified against annual stocking records to determine the hatchery strain and thus lake of origin for each fish. This method was used to collect hatchery–reared Lake Charr originating from Lakes Huron, Lake Champlain, and the humper morph from Lake Superior. Lake Charr morphs from Lake Superior were distinguished using the visual determination methods described in ref^52^. A 1 – 3 gram (g) piece of muscle tissue was collected near the dorsal fin of each fish and frozen on dry ice or preserved in 1 mL DNA/RNA Shield (Zymo Research, Inc.) and sent to the University at Buffalo for storage and further processing.

### Genome Sequencing: Short-Reads

For short-read sequencing, DNA was extracted from 1 mg of muscle tissue from each fish using PureLink Genomic DNA Kit (Invitrogen, Inc.) according to the manufacturer’s protocol. Extracted DNA was sent to Novogene (Davis, CA) for Illumina paired-end sequencing (150 bp x 2) on the NovaSeq 6000 platform until at least 36 Gb (∼15x coverage) of data were generated per sample. We also downloaded moderate coverage short-read data from three lean and three siscowet morph individuals from NCBI (BioSample accessions SAMN39937997 – SAMN39938002 originally published in ref^23^). Illumina data were quality filtered with a Phred score cutoff of 30 (-q 30), and adapter sequences were automatically detected and removed using Trim Galore v0.6.6^152^. Illumina data from a single sample from Lake Champlain was found to have been contaminated with DNA from a second population and was, therefore, removed from further analyses.

### Assessment of Population Structure

Illumina paired-end reads from each sample were mapped to the current *S. namaycush* reference genome (from ref. ^153^; GCF_016432855.1) using bwa-mem2 v2.2.1^154^. Genome-wide single-nucleotide variants (SNVs) were detected using bcftools^155^ v1.15 *mpileup -C50* and *call* using default parameters. To retain only bi-allelic SNVs, remove rare variants (minor allele frequencies < 0.05), and remove all loci with missing data, VCFtools ^156^ v0.1.17 was applied to the SNV data. Population genomic structure within and among sample lakes was visualized using a principal components analysis, which was performed using the glPca function from the adegenet v2.1.3^157^ package in R.

### Long-Term Demographic History

Demographic history was inferred for each sample using the Pairwise Sequentially Markovian Coalescence (PSMC v0.6.5-r67) model^158^. PSMC analyses were run with the parameters established for this species in ref^23^. Briefly, 25 iterations per sample, using parameters -t5 -r5 -p “4+20*2+6*4+4”. These parameters define the maximum time to the most recent common ancestor (t), the ratio of mutation rate to recombination rate (θ/ρ), and the number of free atomic time intervals (p), respectively. A nuclear mutation rate of 7.26e-09 mutations/site/generation and a generation time of 16 years were used to calibrate the PSMC plots via the psmc_plot.pl utility. Effective population size trajectories were visualized using custom R scripts.

### Identifying Runs of Homozygosity

Runs of homozygosity (ROH) and genome-wide heterozygosity were assessed independently for each sample using ROHan^159^ v1.0.1. Initial heterozygosity was calculated in 100 Kb windows. ROH were then identified as 100 Kb windows with heterozygosity below 2.9×10-4 *-rohmu 2.9e-4*. ROHan was then used to apply a Hidden Markov Model to merge adjacent ROH based on a probabilistic framework. Custom R scripts were used to calculate individual the total length of ROH for each sample, and to calculate the average length of ROH within each population.

### Long-Read Sequencing and Genome Assembly

For long-read sequencing, a 1g flash-frozen muscle plug was pulverized in a 5 µL Dounce homogenizer, and high-molecular-weight (HMW) DNA was extracted using a Qiagen Genomic-tip 500/G kit following the manufacturer’s protocol. DNA was quantified on the Qubit 3.0 and checked for protein contamination using a NanoDrop 2000/2000c spectrophotometer. A Short Read Eliminator Kit (Circulomics, Inc.) was used to reduce the number of DNA fragments shorter than 25 kilobases. Size-selected HMW DNA was then used as input in LSK112 ligation preparations and sequenced using one to five R10.4.1 PromethION flow cells until 36 – 133 Gb (∼15-57x genome coverage) of passed reads were produced for moderate-coverage and high-coverage assemblies, respectively. ONT long-reads were basecalled using the super high accuracy basecalling mode in Guppy v5.1.0. A 1g muscle plug from each of the four samples with high-coverage ONT data was sent to Phase Genomics for chromatin conformation capture sequencing of formalin cross-linked DNA on the Illumina NovaSeq 6000 platform.

To maximize the contiguity of the draft genome assemblies for each *S. namaycush* sample, we attempted to optimize the parameters in Flye^65^ v2.9.3 to best match the read depth and sequence length data produced for each sample. Specifically, each genome was assembled at least 3 times, first using all reads >1 Kb in length, with the minimum read overlap setting (*–ovlp*) set to 5 Kb, 7.5 Kb, or 10 Kb for samples with read-length N50s of ≤ 5 Kb, 5.5 - 9 Kb, and >9 Kb, respectively. Next, reads shorter than the read N90 length for each sample were removed using SeqKit v2.9.0^160^, and the remaining reads were reassembled, adjusting to match the overlap-read length rules mentioned above when needed. This process was continued for each sample until the contig length N50 and N90 stopped increasing. The contig-level assembly with the highest N50 and N90 was selected for each assembly and used in the remainder of the genome assembly pipeline. We implemented an iterative polishing approach to correct base errors and small indels inherent in long-read assemblies. Each contig-level assembly was first polished using Medaka v2.0.1(https://github.com/nanoporetech/medaka), followed by three rounds of polishing using SRS data with Pilon^161^ v1.23. We used three iterative rounds of Pilon polishing because the initial correction of widespread errors improves read mapping accuracy in subsequent passes, allowing the tool to resolve increasingly complex error clusters that a single round might miss.

To scaffold high-coverage assemblies, HiC data were mapped to the polished contig-level assemblies using BWA-mem2^154^, and Juicer^162^ v1.6 was used to produce contact heatmaps. Next, contigs were corrected and assembled into chromosomes using 3D-DNA^163^. Finally, the Juicebox^163^ assembly tools were used to visualize and manually edit misassemblies. RagTag^164^ was used to scaffold each moderate-coverage assembly using the high-coverage assembly that was most genetically similar based on a principal components analysis of genome-wide SNV and proportion of shared ancestry an admixture analysis. Finally, scaffolding gaps in all assemblies were closed using TGS-GapCloser^165^ v1.2.0 using ONT long-reads >5 Kb in length. The completeness of each assembly was assessed using BUSCO_v5 ^166^ performed using gVolante2 ^167^ with the Actinopterygii_obd10 database.

### Genome Annotation

Gene models for all 31 *S. namaycush* genomes were produced using the Krabbenhoft Lab Annotation pipeline (https://github.com/KrabbenhoftLab/genome_annotation_pipeline). First, RepeatModeler v2.01^168^ and the “Vertebrata” database from Dfam_3^169^ were used to identify repetitive elements in each assembly. RepeatMasker version v4.1.1^170^ was then used to identify and soft-mask repetitive elements using the custom repeat library generated for each assembly. Gene Model Mapper (GeMoMa) v1.71^171^ was used to generate homology-based gene predictions for each masked assembly using annotations for 12 salmonid species and *Esox lucius* (**Supplementary Table 2**) as “references” to predict gene models using the module *GeMoMaPipeline*. Gene models generated from each reference species were then aggregated and filtered to retain only those with the highest support values using the GeMoMa annotation finalizer (GAF) using default parameters^170^. For each assembly, gene models were also generated with BRAKER3 v3.0.3^172^ using publicly available RNA-seq data from Seneca Lake origin *S. namaycush*^153^ and Lake Superior siscowet and lean morph individuals^45,116^, which were mapped to each assembly using HISAT2^173^, and the same gene annotation data used for GeMoMa predictions as input. Finally, EVidenceModeler v2.0.0^174^ was used to merge GeMoMa and BRAKER3 gene models to produce a final, high-quality genome annotation set for each assembly.

### Detection of Structural Variations

We used the pangenome graph builder pipeline PGGB^68^ v0.5.4 to generate a pangenome graph for *S. namaycush*. To run PGGB, separate graphs were produced for all homologous chromosomes independently, each consisting of a single chromosome from the 31 assemblies. Genome graph files from each chromosome were decomposed and converted into variant call format (vcf) using the vgtoolkit^175^ v1.63. The high-coverage *de novo* assembly from Seneca Lake (SE08) was chosen as the “reference” genome for the vg output to provide a common set of coordinates for the vcf files. Vcf files for each chromosome were then merged and filtered to retain only mutations affecting at least 50 bp (structural variations) using bcftools^155^ v1.20. To validate SVs detected in the pangenome, we used two additional approaches to detect SVs. First, we mapped all ONT data to the SE08 genome using minimap2, then used Sniffles2^70^ v2.2 to identify SVs requiring a minimum of five reads to validate an SV. Second, we mapped all 42 chromosomes from each assembly to the corresponding chromosomes in the SE08 genome using minimap2 and used SVIM-ASM^69^ v1.0.3 to call SVs. Finally, to retain only high-confidence SVs, we used SURVIVOR^176^ v1.0.7 to retain only SVs detected by at least two of the aforementioned methods. If two or more SVs had both start and end positions occurring within 1 Kb of each other, they were merged. Bedtools^177^ 2.31.0 was then used to identify SVs that overlapped genes and repeat sequences. We used GENESPACE v1.3.1^178^ to visualize synteny among the 31 *S. namaycush* assemblies. Gene annotations were first filtered to retain only genes located on one of the 42 chromosomes and the longest transcript per gene using AGAT v0.9.2 ^179^; https://github.com/NBISweden/AGAT). Filtered annotation files were converted to bed files using Bedtools^177^ and protein-peptide FASTA files using AGAT, and used as input for GENESPACE.

### Identifying Early and Late-diverging Ohnologs

Ohnolog pairs within the *Salvelinus namaycush* genome were identified using a self-self synteny analysis using the following tools on CoGe^180^ (Comparative Genomics, genomevolution.org). SynMap2^181^ was used to perform BLASTn searches between all genes in one assembly per population. The DAGchainer algorithm^181^ was implemented to identify synteny blocks, with a maximum gap of 5 genes (-D) between gene pairs and a minimum of 10 aligned gene pairs required to call a syntenic block. CodeML from the PAML4^182^ package was used to calculate synonymous substitution per site values (*K_s_*) between each gene pair. Ohnologs were identified as gene pairs occurring in regions of conserved synteny between two different chromosomes. Late-diverging regions were identified as homeologous syntenic blocks in which at least 30% of genes in that block had ≥90% sequence similarity and a log10(*K_s_*) < 0.99 with their Ohnologs. Late-diverging Ohnologs were defined as all genes within a late-diverging syntenic block that retained a gene pair on their homeologous chromosomes. All other homeologous syntenic blocks and the Ohnologs contained within them were defined as late-diverging regions and late-diverging Ohnologs, respectively. To avoid complications arising from Ohnologs remaining from the Teleost-Specific Third Round whole genome duplication (3R), Ohnolog pairs were filtered to retain only their reciprocally best BLAST hits.

### Calling CpG DNA Methylation

Raw Nanopore sequencing data for the high-coverage lean and siscowet samples were rebasecalled using the algorithm Guppy v6.4.8+31becc9, with the dna_r10.4_e8.1_modbases_5hmc_5mc_cg_sup_prom.cfg model used to perform nucleotide basecalling and direct DNA methylation detection. This model was chosen to allow for “super-accurate” (SUP) basecalling (mean basecalling accuracy of 99.0%) as well as the identification of two types of DNA methylation, 5-methylcytosine (CpGm), and 5-hydroxymethylcytosine. To call modified bases, all reads from each sample were first mapped to their own chromosome-level genome assembly using minimap2 to generate a modBAM file. Next Modkit (v0.3.0) pileup with the *–cpg* flag was used to convert modBAM files into bedMethyl files containing genome-wide DNA methylation calls at all CpG dinucleotide sites for each sample. Bedtools^177^ 2.31.0 was then used to identify CpGm sites that overlapped genes using a bedfile version of each assemblies’ annotation file.

### Producing Haplotype-Resolved Assemblies

To produce haplotype-resolved assemblies for lean and siscowet *S. namaycush*, ONT reads were error-corrected using the deep-learning-based haplotype-aware read correction tool PECAT^183^ v0.0.3. Corrected long-reads and HiC data were then used as inputs in Hifiasm^184^ v0.19.6 to produce a draft phased assembly. To avoid known issues with contained read exclusion in Hifiasm^184^, corrected reads >30 Kb in length were trimmed or split using SeqKit^160^ v2.9.0. To scaffold the assembled contigs of each haplotype into chromosomes, Hi-C data were first mapped to each contig-level assembly using bwa-mem2 v2.2 ^154^ and scaffolded using YaHS v1.2.2 ^185^. Finally, Juicebox ^163^ was used to visualize and manually edit misassemblies.

## STATISTICS AND REPRODUCIBILITY

### DATA AVAILABILITY

The data underlying this article are available in the NCBI Sequence Read Archive (SRA) and GenBank databases. The raw long-read (ONT), short-read (Illumina), and Hi-C sequencing data have been deposited in the SRA under the BioProject PRJNA1453007 under the Biosample accessions SAMN57248128 - SAMN57248158. The 31 chromosome-level genome assemblies generated in this study have been deposited in GenBank with the following genome accessions: JBXUKS000000000, JBXUKR000000000, JBXUKQ000000000, JBXUKP000000000, JBXUKO000000000, JBXUKN000000000, JBXUKM000000000, JBXUKL000000000, JBXUKK000000000, JBXUKJ000000000, JBXUKI000000000, JBXUKH000000000, JBXUKG000000000, JBXUKF000000000, JBXUKE000000000, JBXUKD000000000, JBXUKC000000000, JBXUKB000000000, JBXUKA000000000, JBXUJZ000000000, JBXUJY000000000, JBXUJX000000000, JBXUJW000000000, JBXUJV000000000, JBXUJU000000000, JBXUJT000000000, JBXUJS000000000, JBXUJR000000000, JBXUJQ000000000, JBXUJP000000000, JBXUJO000000000. Custom scripts used for genome assembly, scaffolding, and downstream analyses are available on GitHub at Github.com/organizations/KrabbenhoftLab/Lake_Charr_Pangenome. Genome annotations, repeat bed files, and protein FASTA files are available on Dryad http//##########.

## ACKNOWLEDGEMENTS

We would like to thank Thomas Detmer from Cornell University’s Department of Natural Resources and the Environment, Ian Harding and the Red Cliff Band of the Lake Superior Chippewa Nation, the crew members of the RV Kaho at the U.S. Geological Survey (USGS) Lake Ontario Biological Station, and Jacques Rinchard at SUNY Brockport for providing samples. We thank Charlotte Lindqvist, Omer Gokcumen, and members of the T. Krabbenhoft Lab for providing helpful feedback on this research. We also thank Nick Sard, Amanda Ackiss, and Brian O’Malley for providing internal reviews of this manuscript to meet USGS publishing standards. Funding for this research was provided by the U.S. Fish and Wildlife Service, the Great Lakes Restoration Initiative, the Great Lakes Research Consortium Small Grants Program, and laboratory startup funds to TJ Krabbenhoft by the University at Buffalo. Any use of trade, product, or firm names is for descriptive purposes only and does not imply endorsement by the U.S. Government. This work made use of the resources of the Center for Computational Research (CCR) at the University at Buffalo, http://hdl.handle.net/10477/79221. The Lake Charr illustrations used in figures 2, 4, and 5 are the work of Paul Vecsei and were used with the permission of the Great Lakes Fishery Commission.

## AUTHOR CONTRIBUTIONS

Conceptualization: CAO, DG, TJK

Methodology: CAO, TJK, NJCB

Software: CAO, NJCB, DJM, SJF

Investigation: CAO

Formal Analysis: CAO, SJF

Resources: BFL, DG, TJK

Data Curation: CAO

Writing – original draft: CAO, TJK

Writing – review & editing: CAO, NJCB, DJM, SJF, BFL, VA, DG, TJK

Visualization: CAO

Supervision: TJK

Project Administration: CAO, TJK

Funding Acquisition: CAO, DG, TJK

## ETHICS STATMENT

All sampling and handling of fish during research are carried out in accordance with guidelines for the care and use of fishes by the American Fisheries Society (http://fisheries.org/docs/wp/Guidelines-for-Use-of-Fishes.pdf).

## SUPPLEMENTARY TABLES AND FIGURES

**Supplementary Table 1.** A list of samples used for moderate-coverage genome assembly, and the respective high-coverage assemblies used as references for scaffolding.

| Sample ID<br>(moderate coverage) | Population | Reference ID<br>(high-coverage) | Reference Population |
| --- | --- | --- | --- |
| AM02 | Little Moose Lake | AM03 | Little Moose Lake |
| AM04 | Little Moose Lake | AM03 | Little Moose Lake |
| FL03 | Farrow Lake | SL01 | Lake Superior: lean |
| FL04 | Farrow Lake | SL01 | Lake Superior: lean |
| FL05 | Farrow Lake | SL01 | Lake Superior: lean |
| HP30 | Lake Huron | SS02 | Lake Superior: siscowet |
| HP46 | Lake Huron | SS02 | Lake Superior: siscowet |
| HP49 | Lake Huron | SS02 | Lake Superior: siscowet |
| KE04 | Keuka Lake | SE08 | Seneca Lake |
| KE15 | Keuka Lake | SE08 | Seneca Lake |
| KE17 | Keuka Lake | SE08 | Seneca Lake |
| LC36 | Lake Champlain | SE08 | Seneca Lake |
| LC47 | Lake Champlain | SE08 | Seneca Lake |
| LC48 | Lake Champlain | SE08 | Seneca Lake |
| SH29 | Lake Superior: humper | SS02 | Lake Superior: siscowet |
| SH45 | Lake Superior: humper | SS02 | Lake Superior: siscowet |
| SK01 | Skaneateles | SE08 | Seneca Lake |
| SK06 | Skaneateles | SE08 | Seneca Lake |
| SK10 | Skaneateles | SE08 | Seneca Lake |
| SL01-21 | Lake Superior: lean | SL01 | Lake Superior: lean |
| SL02 | Lake Superior: lean | SL01 | Lake Superior: lean |
| SS03 | Lake Superior: siscowet | SS02 | Lake Superior: siscowet |
| SS04 | Lake Superior: siscowet | SS02 | Lake Superior: siscowet |
| SE03 | Seneca Lake | SE08 | Seneca Lake |
| SE07 | Seneca Lake | SE08 | Seneca Lake |
| SE09 | Seneca Lake | SE08 | Seneca Lake |
| SE16 | Seneca Lake | SE08 | Seneca Lake |

**Supplementary Table 2.** Genome assemblies used as references for homology-based gene predictions for the 31 *Salvelinus namaycush* assemblies.

| **Common Name (Species)** | **NCBI Accession** |
| --- | --- |
| Lake Whitefish (*Coregonus clupeaformis*) | GCF_020615455.1 |
| Atlantic Salmon (*Salmo salar*) | GCF_905237065.1 |
| Alpine Whitefish (*Coregonus* sp. 'balchen') | GCA_902810595.1 |
| Rainbow Trout (*Oncorhynchus mykiss*) | GCF_013265735.2 |
| Lake Charr (*Salvelinus namaycush*) | GCF_016432855.1 |
| Brown Trout (*Salmo trutta*) | GCF_901001165.1 |
| Chinook Salmon (*Oncorhynchus tshawytscha*) | GCF_018296145.1 |
| Chum Salmon (*Oncorhynchus keta*) | GCF_023373465.1 |
| Coho Salmon (*Oncorhynchus kisutch*) | GCF_002021735.2 |
| Pink Salmon (*Oncorhynchus gorbuscha*) | GCF_021184085.1 |
| Sockeye Salmon (*Oncorhynchus nerka*) | GCF_006149115.2 |
| Arctic Charr (*Salvelinus* sp.) | GCF_002910315.2 |
| Northern Pike (*Esox lucius*) | GCF_011004845.1 |

**Supplementary Table 3.** Genome annotation BUSCO scores for 31 *Salvelinus namaycush* assemblies.

| Sample ID | Lake | Percent of BUSCO Actinopterygii_db10 (n=3640) |  |  |  |  |
| --- | --- | --- | --- | --- | --- | --- |
|  |  | Single | Duplicate | Fragmented | Missing | Complete |
| AM02 | Little Moose | 58.4 | 36.4 | 2.4 | 2.8 | 94.8 |
| AM03 | Little Moose | 58.0 | 37.8 | 2.0 | 2.2 | 95.8 |
| AM04 | Little Moose | 58.9 | 36.1 | 2.2 | 2.8 | 95.0 |
| FL03 | Farrow | 58.2 | 37.6 | 1.8 | 2.4 | 95.8 |
| FL04 | Farrow | 57.9 | 37.6 | 2.0 | 2.5 | 95.5 |
| FL05 | Farrow | 58.3 | 37.3 | 2.0 | 2.4 | 95.6 |
| HP30 | Lake Huron | 57.6 | 38.2 | 1.7 | 2.5 | 95.8 |
| HP46 | Lake Huron | 57.2 | 38.5 | 2.0 | 2.3 | 95.7 |
| HP49 | Lake Huron | 57.1 | 38.6 | 1.9 | 2.4 | 95.7 |
| KE04 | Keuka Lake | 57.0 | 37.3 | 2.4 | 3.3 | 94.3 |
| KE15 | Keuka Lake | 57.2 | 38.2 | 2.0 | 2.6 | 95.4 |
| KE17 | Keuka Lake | 58.9 | 36.5 | 1.9 | 2.7 | 95.4 |
| LC36 | Lake Champlain | 58.0 | 37.3 | 2.1 | 2.6 | 95.3 |
| LC47 | Lake Champlain | 58.3 | 37.3 | 2.0 | 2.4 | 95.6 |
| LC48 | Lake Champlain | 57.3 | 38.7 | 1.8 | 2.2 | 96.0 |
| SE03 | Seneca Lake | 58 | 37.9 | 1.9 | 2.2 | 95.9 |
| SE07 | Seneca Lake | 57.8 | 38.0 | 1.9 | 2.3 | 95.8 |
| SE08 | Seneca Lake | 57.4 | 38.3 | 2.0 | 2.3 | 95.7 |
| SE09 | Seneca Lake | 58.3 | 37.6 | 1.9 | 2.2 | 95.9 |
| SE16 | Seneca Lake | 57.3 | 38.4 | 1.7 | 2.6 | 95.7 |
| SH29 | Lake Superior:<br>humper | 57.4 | 38.0 | 2.0 | 2.6 | 95.4 |
| SH45 | Lake Superior:<br>humper | 57.4 | 37.4 | 2.0 | 3.2 | 94.8 |
| SK01 | Skaneateles Lake | 57.3 | 38.2 | 2.0 | 2.5 | 95.5 |
| SK06 | Skaneateles Lake | 57.7 | 37.9 | 2.0 | 2.4 | 95.6 |
| SK10 | Skaneateles Lake | 56.5 | 39.4 | 2.0 | 2.1 | 95.9 |
| SL01 | Lake Superior: lean | 57.4 | 38.3 | 2.0 | 2.3 | 95.7 |
| SL01-21 | Lake Superior: lean | 58.4 | 37.0 | 2.2 | 2.4 | 95.4 |
| SL02 | Lake Superior: lean | 58.1 | 36.6 | 2.4 | 2.9 | 94.7 |
| SS02 | Lake Superior:<br>siscowet | 56.8 | 38.5 | 2.2 | 2.5 | 95.3 |
| SS03 | Lake Superior:<br>siscowet | 57.5 | 38.3 | 1.8 | 2.4 | 95.8 |
| SS04 | Lake Superior:<br>siscowet | 57.2 | 38.3 | 1.9 | 2.6 | 95.5 |

**Supplementary Table 4.** Genes contained within the boundaries of the putative 1 – 3 translocation in siscowet and humper *Salvelinus namaycush*.

| SS02 Gene ID | KEGG Ortholog Number | Gene Name |
| --- | --- | --- |
| evm.model.chr3.1220 | K21436 | *ankrd11*; ankyrin repeat domain-containing protein 11 |
| evm.model.chr3.1221 |  | *slc22a31* |
| evm.model.chr3.1222 | K01273 | *dpep*; membrane dipeptidase [EC:3.4.13.19] |
| evm.model.chr3.1223 |  | *sult2b1* |
| evm.model.chr3.1224 | K16142 | *hp*; haptoglobin |
| evm.model.chr3.1225 |  | *taar13c* |
| evm.model.chr3.1226 | K00254 | *dhodh*, pyrD; dihydroorotate dehydrogenase [EC:1.3.5.2] |
| evm.model.chr3.1227 |  | PKD1L2 |
| evm.model.chr3.1228 | K19476 | IST1; vacuolar protein sorting-associated protein IST1 |
| evm.model.chr3.1229 | K14559 | MPP10; U3 small nucleolar RNA-associated protein MPP10 |
| evm.model.chr3.1230 | K05606 | MCEE, epi; methylmalonyl-CoA/ethylmalonyl-CoA epimerase [EC:5.1.99.1] |
| evm.model.chr3.1231 | K20055 | APBA2, X11L; amyloid beta A4 precursor protein-binding family A member 2 |
| evm.model.chr3.1232 | K00008 | SORD, gutB; L-iditol 2-dehydrogenase [EC:1.1.1.14] |
| evm.model.chr3.1233 | K25750 | *terb2*; telomere repeats-binding bouquet formation protein 2 |
| evm.model.chr3.1234 | K05701 | TJP1, ZO1; tight junction protein 1 |
| evm.model.chr3.1235 |  | *cerkl*; ceramide kinase-like |
| evm.model.chr3.1236 |  | *fam189a1* |
| evm.model.chr3.1237 |  | *golm2*; Golgi membrane protein 2 |
| evm.model.chr3.1238 | K17616 | CTDSPL2; CTD small phosphatase-like protein 2 [EC:3.1.3.-] |
| evm.model.chr3.1239 | K24636 | *strc*; stereocilin |
| evm.model.chr3.1240 | K03245 | EIF3J; translation initiation factor 3 subunit J |
| evm.model.chr3.1241 | K24436 | CILP; cartilage intermediate layer protein |
| evm.model.chr3.1242 |  |  |
| evm.model.chr3.1243 |  | ZDBF2; Zinc Finger DBF-Type Containing 2 |
| evm.model.chr3.1244 | K22654 | IGDCC3, PUNK; immunoglobulin superfamily DCC subclass member 3 |

**Fig. S1.**
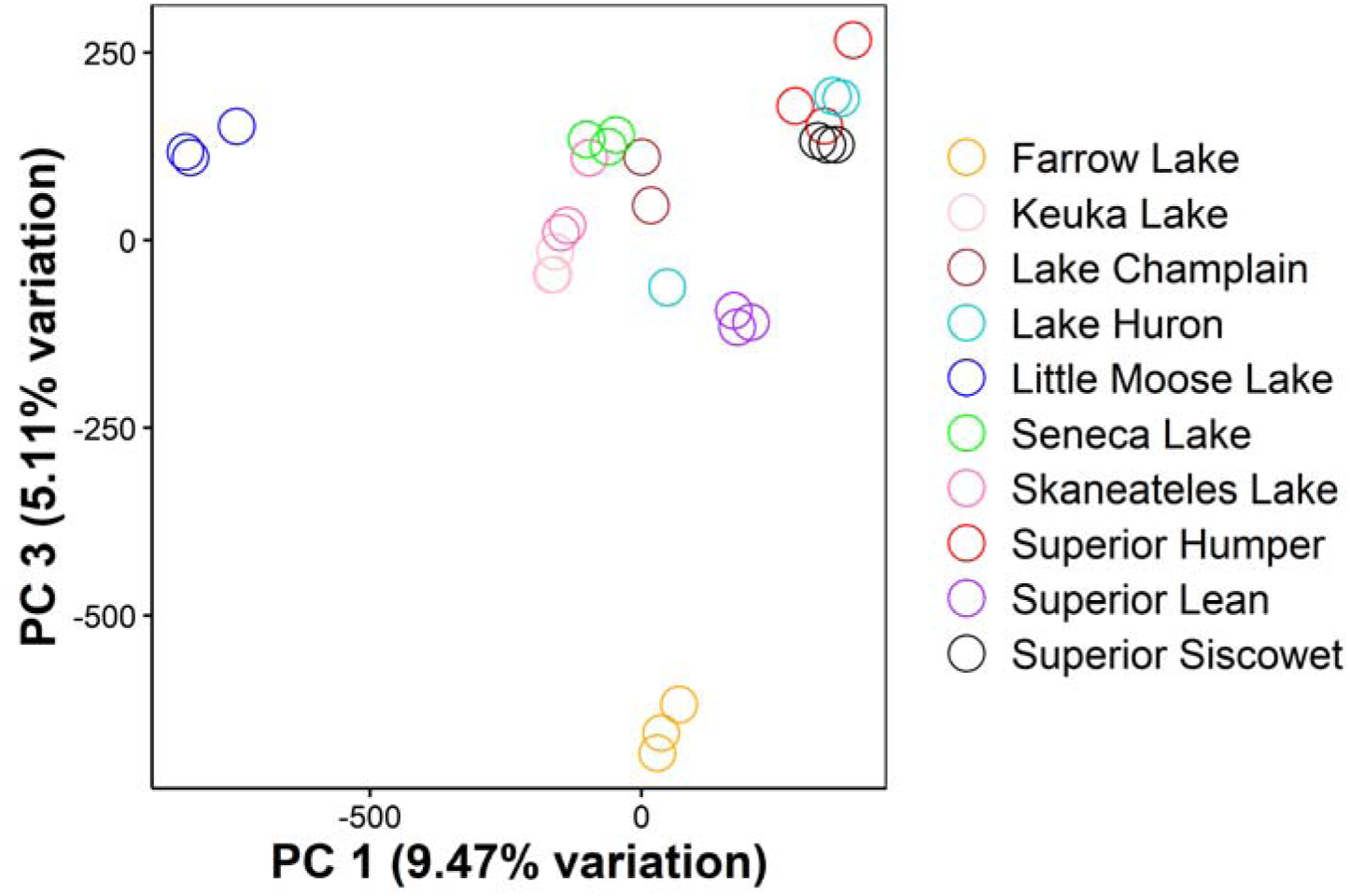
A principal components analysis using genome-wide single-nucleotide variants detected in 10 populations of *Salvelinus namaycush*. The x- and y-axes depict variation across principal components (PC) 1 and 3, respectively. The Farrow Lake population, although geographically closest to the Little Moose Lake population, exhibits considerable divergence along PC3. The color of each point corresponds to the lake of origin for each sample.

**Fig. S2.**
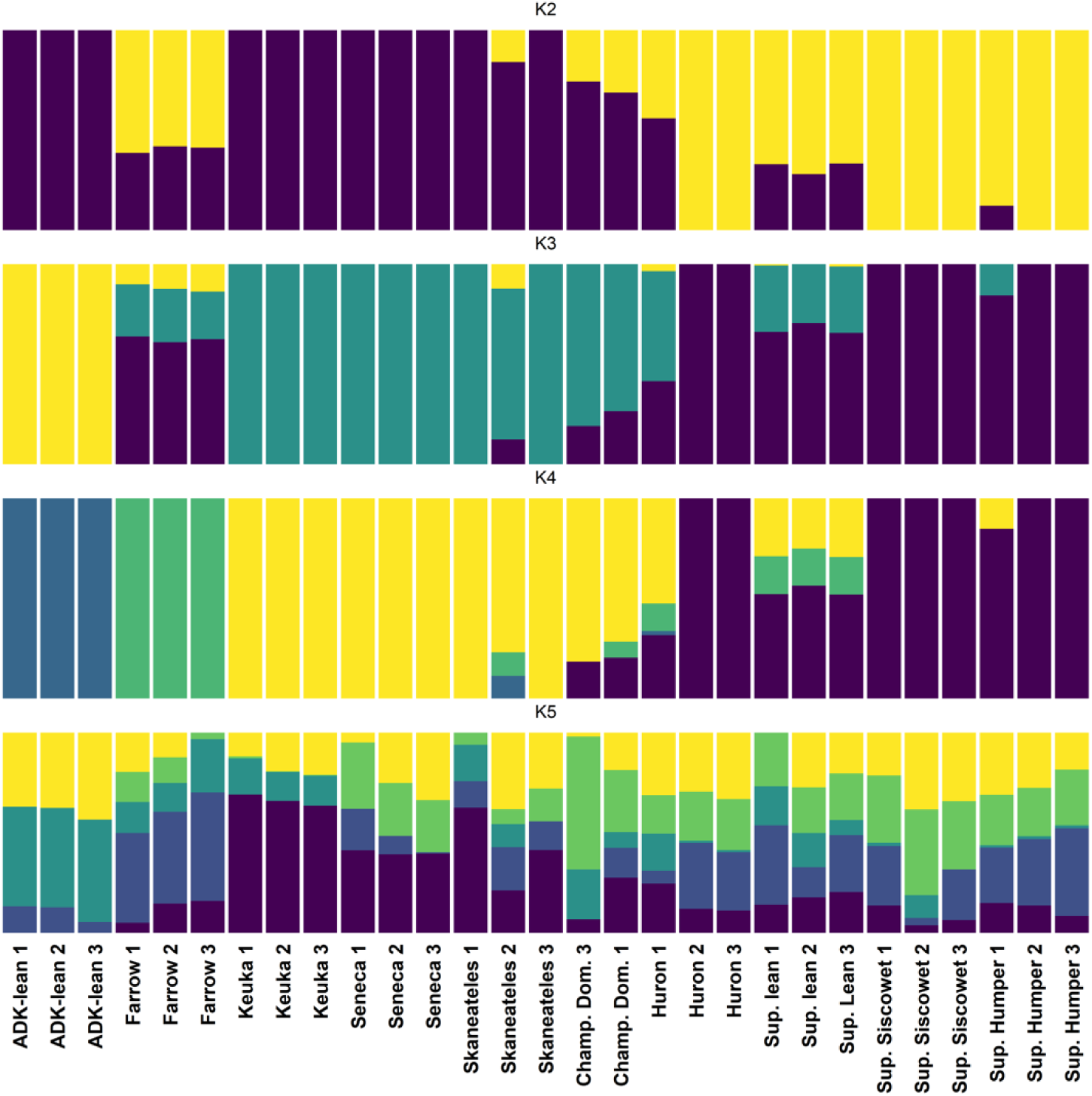
Admixture analysis of single-nucleotide variants performed using ADMIXTURE^186^ displaying models for 2 – 5 ancestral groups (K2 – K5).

**Fig. S3.**
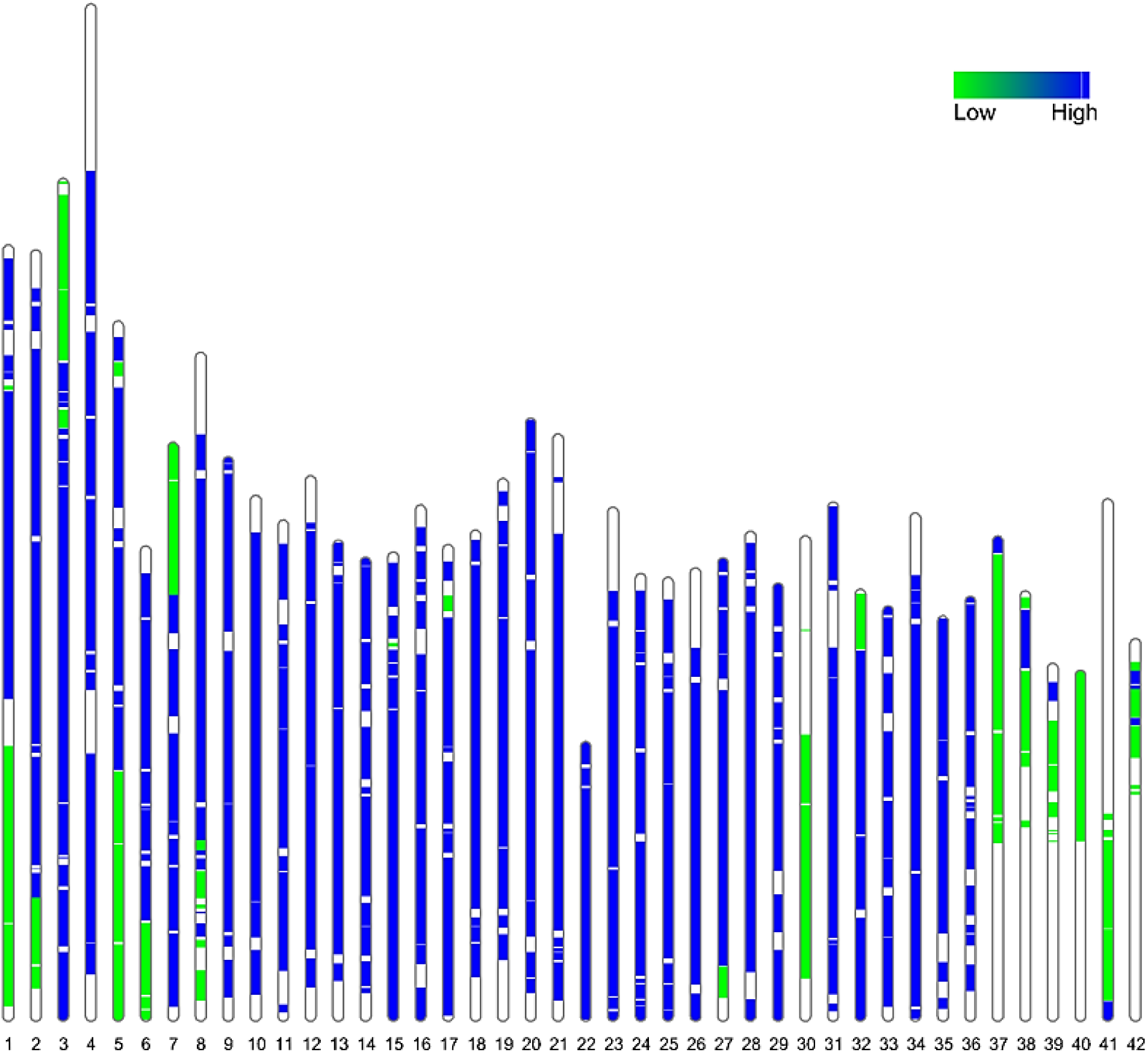
Ideogram displaying the genomic location of early- and late-diverging regions on the 42 *Salvelinus namaycush* chromosomes. Each chromosome is painted according to the level of sequence divergence between it and its homeologous region. High-divergence regions (i.e., those containing early-diverging Ohnologs) are colored blue, and low-divergence regions (i.e., those containing late-diverging Ohnologs) are colored green. Chromosome segments with no color were found to have little to no synteny with any other regions of the genome.

**Fig. S4.**
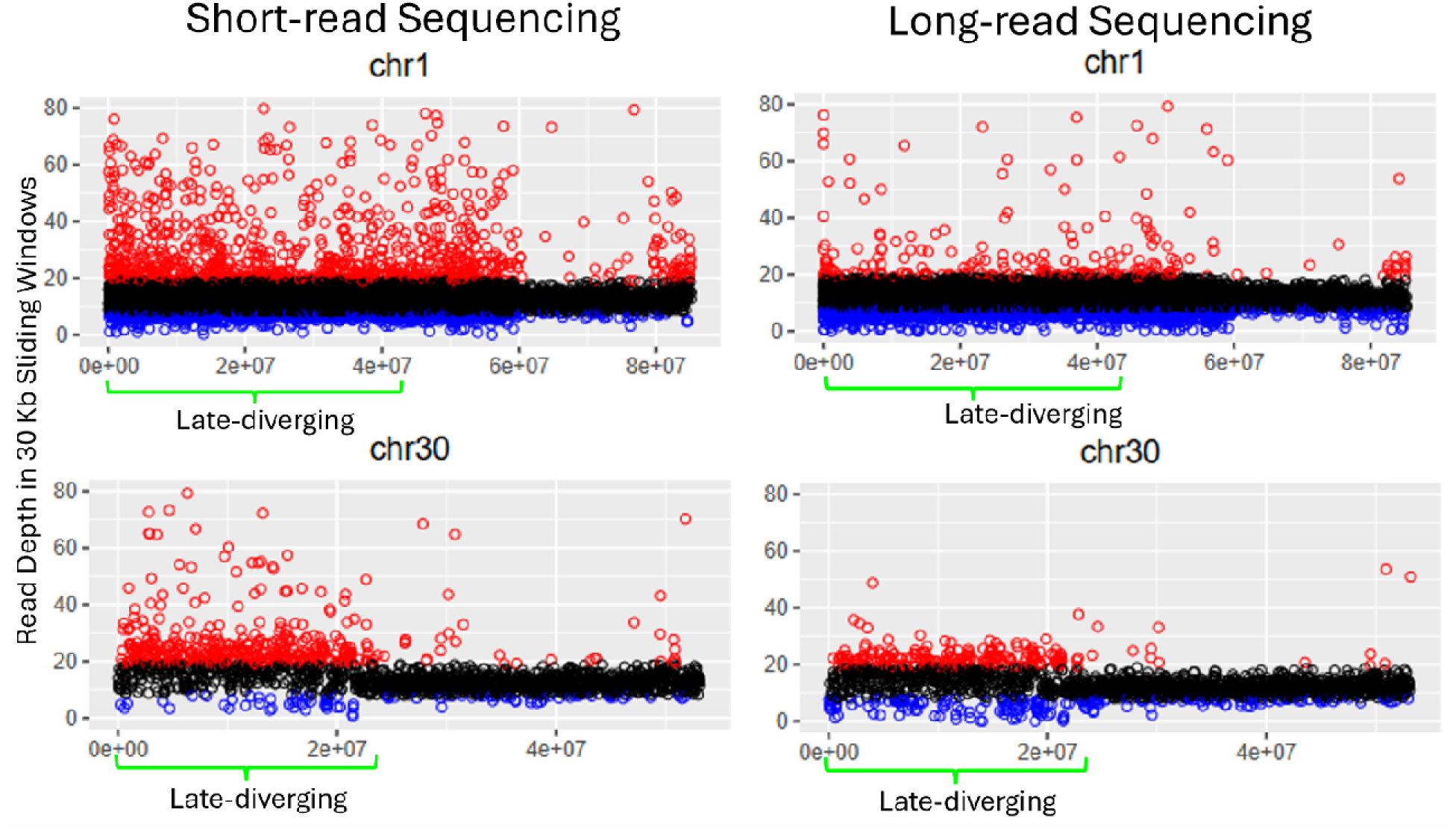
Read depth of short- and long-read sequencing data mapped to chromosomes 1 and 30 of *Salvelinus namaycush*. Read depth was calculated in 30 Kb sliding windows. Points are colored according to their relationship to the genome-wide average read depth; red points represent windows greater than or equal to 1.5x the genome-wide average, blue dots are those with less than or equal to 0.5x, and black dots are all other windows. Short reads have artificially inflated read depth in late-diverging regions, while this problem is greatly reduced in long reads.

**Fig. S5.**
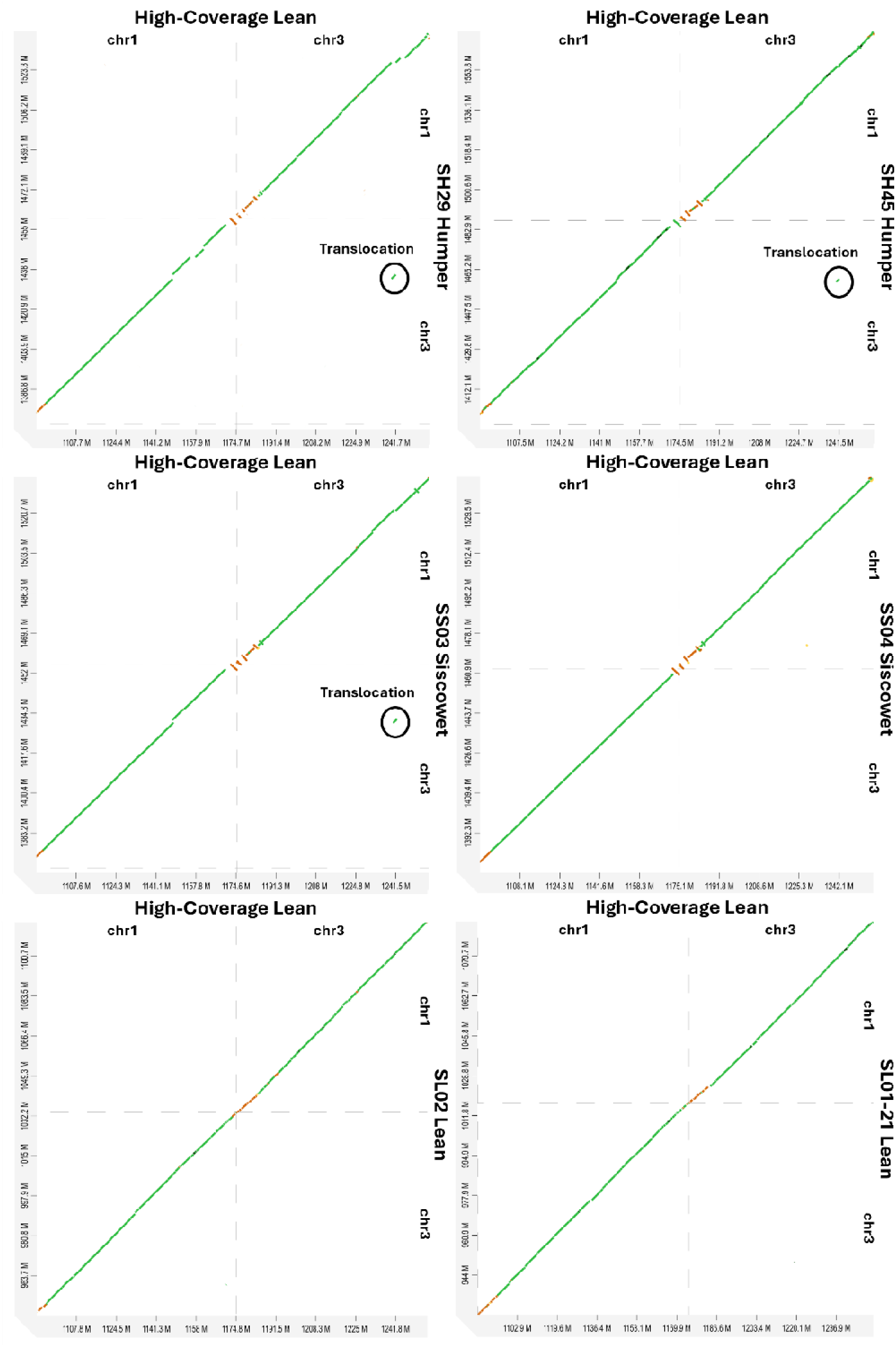
Dot plots showing the putative chromosome 1-to-3 translocation in moderate coverage siscowet and humper genome assemblies, and its absence from the moderate coverage lean assemblies. Each pair of moderate-coverage chromosomes was mapped to chromosomes in the high-coverage assembly of the Lake Superior lean morph. Dot plots were generated using D-GENIES^187^ using minimap2 for mapping.

**Fig. S6.**
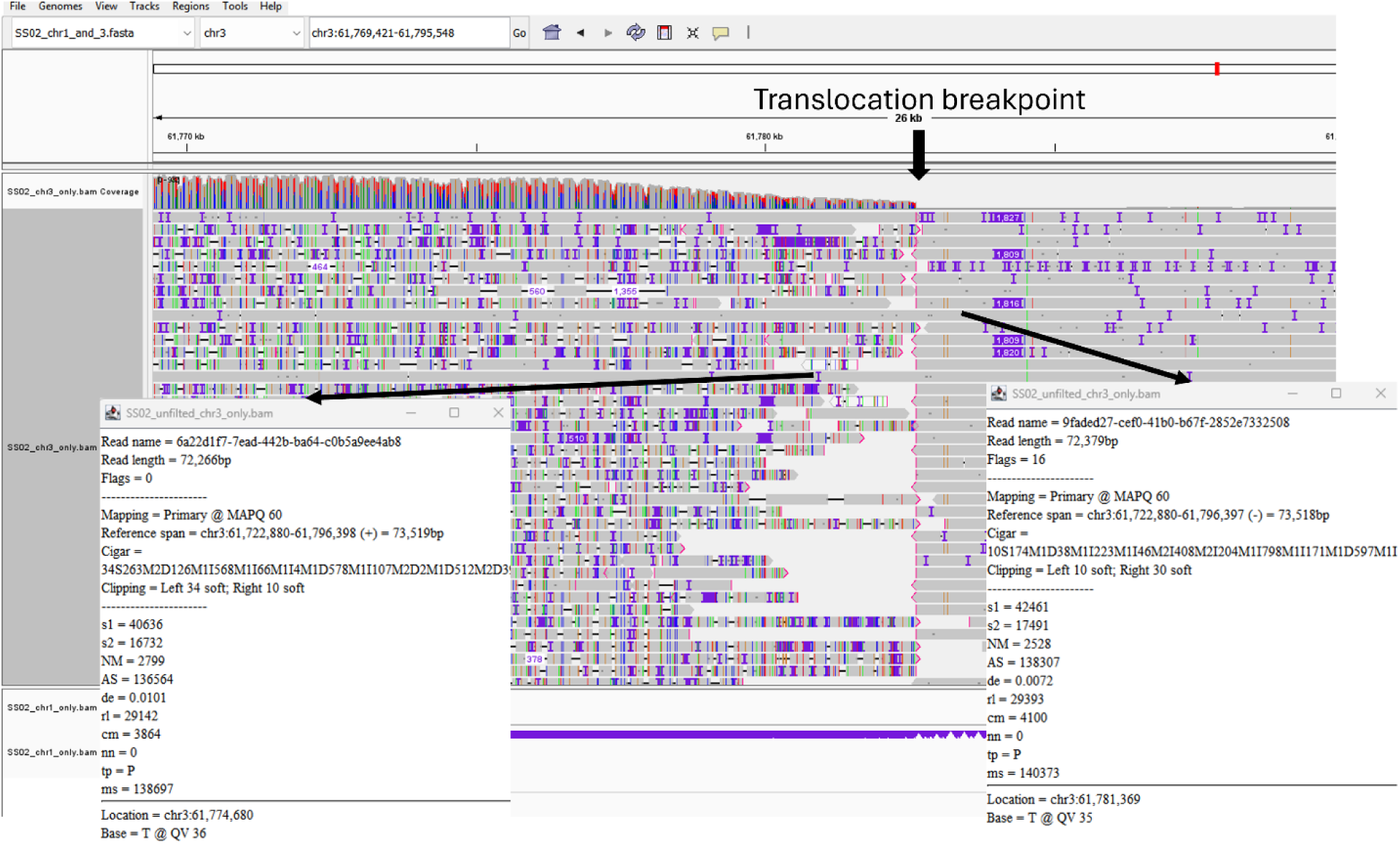
Long-read alignment diagnostics at the chromosome 3 translocation breakpoint highlighting a potential assembly artifact mediated by an ultra-long duplex read. Integrated Genomics Viewer (IGV) snapshot of Oxford Nanopore Technologies (ONT) long reads mapped back to the high-coverage siscowet haplotype assembly around the putatively translocated breakpoint region on chromosome 3. The standard long reads terminate with soft-clipping (indicated by vertical pink boundaries) at the translocation breakpoint. The continuous, unclipped path through this breakpoint is supported exclusively by two ultra-long (UL) reads (detailed in the lower-left and lower-right callouts). Notably, these two molecules map to opposite strands (positive and negative orientations, respectively), share near-identical sequence lengths (72,266 bp and 72,379 bp), and align to virtually identical genomic coordinates (chr3:61,722,880–61,796,398). This combination of symmetric features provides definitive molecular evidence of an ONT read duplex - where the template and complement strands of a single, physical *in vitro* chimeric molecule were sequenced sequentially through a single pore.

**Fig. S7.**
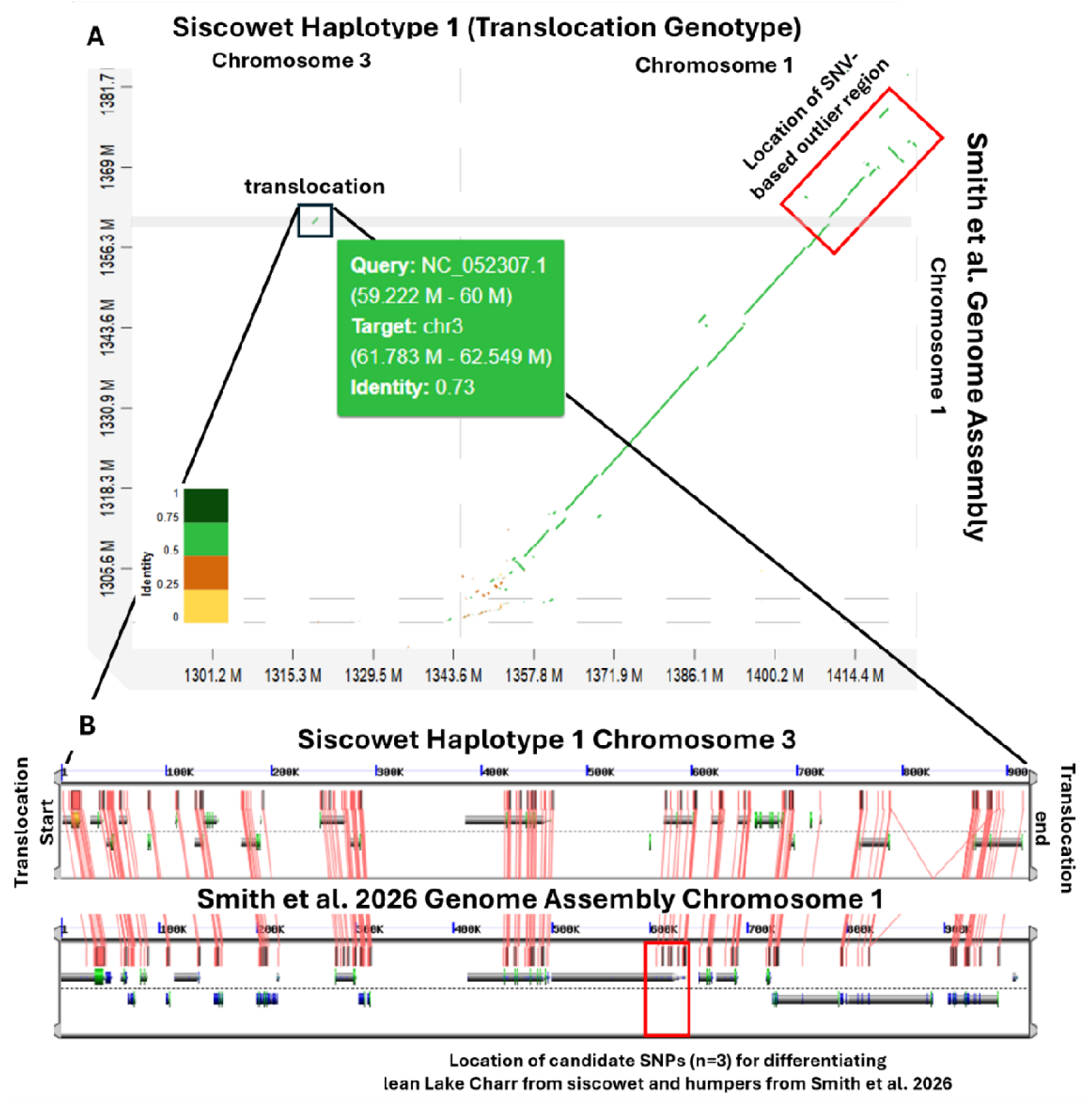
**A**, Dot plot showing the putative chromosome 1-to-3 translocation between the high-coverage siscowet assembly produced in this study and the genome assembly used as a reference for a RAD-seq analysis designed to identify genomic regions of divergence between lean, siscowet, and humper morphs (ref ^123^). The putative translocation is bounded by a black box, and the boundaries are highlighted with a grey box. The location of the largest outlier peak identified in ref^123^ is bounded by a red box. The dot plot was generated using D-GENIES^187^ using minimap2 for mapping. **B**, Synteny and gene structure of the genes found in this region in siscowet and their syntelogs on chromosome 1 in the reference genome used by Smith et al. The red box depicts the location of the three outlier SNVs found to distinguish lean from siscowet and humper morphs. The Genome Evolution Analysis tool from CoGe^180^ (Comparative Genomics, genomevolution.org) was used to create and visualize these results. Exons are depicted as green polygons and introns are shown as grey bars. Pink lines depict the best-matching coding sequences for each exon between assemblies, based on BLASTn results.

**Fig. S8.**
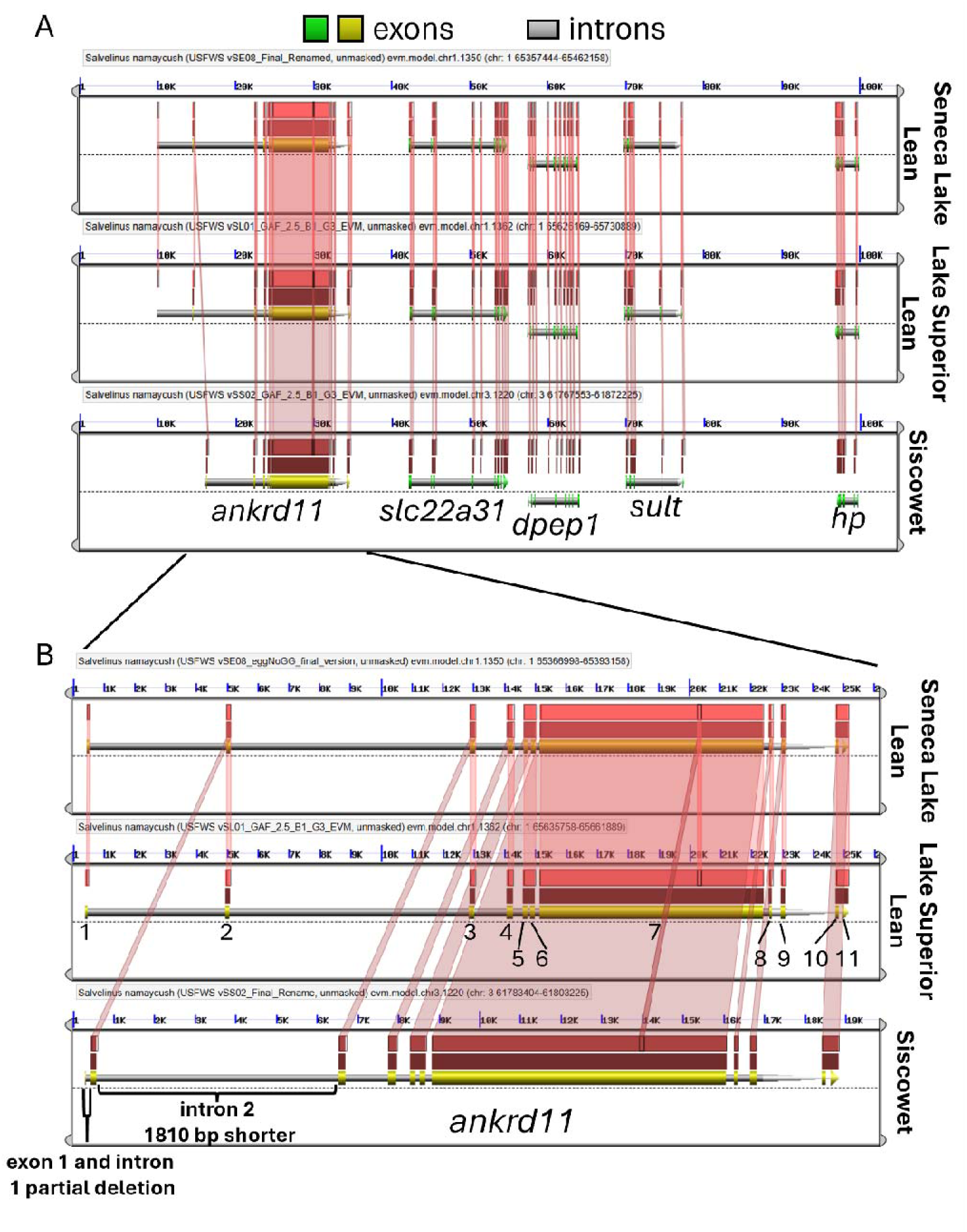
**A**, Synteny and gene structure of the first five genes found on chromosome 1 in lean morph *Salvelinus namaycush* and on the homologous region translocated to chromosome 3 in siscowet. **B**, Gene structure and sequence alignment results for *ankyrin repeat domain containing protein 11* (*ankrd11*) showing a partial deletion of the first exon and intron, and another partial deletion of the second intron in siscowet. The Genome Evolution Analysis tool from CoGe^180^ (Comparative Genomics, genomevolution.org) was used to create and visualize these results. Exons are depicted as yellow and green polygons for *ankrd11* and the other four genes, respectively. Introns are shown as grey bars. Pink and red lines depict the best-matching coding sequences for each exon between assemblies, based on BLASTn results.

